# The study of the determinants controlling Arpp19 phosphatase-inhibitory activity reveals a new Arpp19/PP2A-B55 feedback loop

**DOI:** 10.1101/2020.05.11.087890

**Authors:** Jean Claude Labbé, Suzanne Vigneron, Francisca Méchali, Perle Robert, Cindy Genoud, Perrine Goguet-Rubio, Phillipe Barthe, Gilles Labesse, Martin Cohen-Gonsaud, Anna Castro, Thierry Lorca

## Abstract

Arpp19 is a potent inhibitor of PP2A-B55 that regulates this phosphatase to ensure the stable phosphorylation of mitotic/meiotic substrates. At G2-M, Arpp19 is phosphorylated by Greatwall on S67. This phosphorylated Arpp19 form displays a high affinity to PP2A-B55 and a slow dephosphorylation rate, acting as an “unfair” competitor of PP2A-B55 substrates. The molecular determinants conferring slow dephosphorylation kinetics to S67 are unknown. PKA also phosphorylates Arpp19. This phosphorylation performed on S109 is essential to maintain prophase I-arrest in Xenopus oocytes although the underlying signaling mechanism is elusive. Here, we characterized the molecular determinants conferring slow dephosphorylation to S67 and controlling PP2A-B55 inhibitory activity of Arpp19. Moreover, we showed that phospho-S109 restricts S67 phosphorylation by increasing its catalysis by PP2A-B55. Finally, we discovered a double feed-back loop between these two phospho-sites which is essential to coordinate the temporal pattern of Arpp19-dependent PP2A-B55 inhibition and Cyclin B/Cdk1 activation during cell division.

## INTRODUCTION

Entry and exit of mitosis and meiosis is induced by oscillations of protein phosphorylation/dephosphorylation. These oscillations are the result of the activation and inactivation of the master kinase Cyclin B/Cdk1 and of its counterbalancing phosphatase PP2A-B55^1–7^. Although substrate phosphorylation is triggered at mitotic entry by the activation of Cyclin B/Cdk1, it is only fully achieved if PP2A-B55 activity is negatively regulated ^8–11^. This regulation is in charge of Arpp19 and ENSA, two members of the endosulfine family of proteins recently identified as two potent inhibitors of PP2A-B55^3,4^. PP2A-B55 inhibition does not only control mitosis but also other phases of the cell cycle. In this line, ENSA has been involved in the negative regulation of this phosphatase during DNA replication by controlling the dephosphorylation and degradation of the replication factor Treslin ^12^. Conversely, Arpp19 has been shown to be an essential gene controlling PP2A-B55 during mitotic division. Indeed, the ablation of this protein in Mouse Embryonic Fibroblasts (MEFs) promotes the premature dephosphorylation of mitotic substrates resulting in the disruption of the correct temporal order of cellular events during mitotic progression ^6^. Both Arpp19 and ENSA are the unique substrates of the kinase Greatwall (Gwl). At G2-M onset, Gwl is activated and phosphorylates Arpp19 at a single site, S67. Arpp19 phosphorylation then triggers its binding and the subsequent inhibition of PP2A-B55, hence allowing the stable phosphorylation of Cyclin B/Cdk1 substrates and mitotic entry ^5,13^. At mitotic exit, Gwl is inactivated resulting in Arpp19 dephosphorylation, PP2A-B55 reactivation and the gradual dephosphorylation of mitotic substrates ^14,15^. The mechanisms by which Arpp19/ENSA inhibit PP2A-B55 are still elusive, however, a previous report demonstrated that ENSA acts as an “unfair” substrate of PP2A-B55 ^16^. Accordingly, data established that S67 phosphorylated ENSA displays a high affinity for this phosphatase but a slow dephosphorylation rate and consequently acts as a major “unfair” competitor of PP2A-B55 substrates. The molecular determinants of Arpp19 and ENSA conferring these specific properties to phospho-S67 are completely unknown.

Besides Gwl-dependent phosphorylation, Arpp19 is also phosphorylated by PKA. PKA-dependent phosphorylation of the alternative splice variant of Arpp19, the protein Arpp16, was firstly described. Arpp16 is enriched in striatal neurons and it is phosphorylated in these cells by the MAST3 kinase. This phosphorylation promotes Arpp16 binding and inhibition of PP2A-B55, a regulation that is essential for striatal signaling ^17^. It has been shown that the additional PKA-dependent phosphorylation of Arpp16 in a second site negatively modulates its inhibitory activity towards PP2A-B55. Besides Arpp16, Arpp19 is also phosphorylated by PKA in Xenopus oocytes. PKA-dependent phosphorylation of Arpp19 on residue S109 is essential to arrest Xenopus oocytes in prophase I of meiosis ^18^. Accordingly, the injection of a S109D phospho-mimetic Arpp19 form to these oocytes blocks meiotic resumption induced by progesterone. Conversely, when Arpp19 is stably phosphorylated “*in vitro”* on S67 by ATP^γs^ and injected to prophase I-arrested oocytes, meiosis is resumed even following overexpression of phospho-S109-Arpp19. This inhibitory effect of phospho-S109 has been attributed to its ability in preventing the formation of a starter amount of active Cdk1 necessary for triggering meiosis. This starter amount would require Gwl activation, S67 Arpp19 phosphorylation and the inhibition of PP2A-B55 an essential process to promote stable protein phosphorylation and meiotic progression ^19,20^. How phospho-S109 Arpp19 negatively regulates the starter amount activation of Cyclin B/Cdk1 is completely unknown. A putative hypothesis was that S109 phosphorylation could regulate Gwl-dependent phosphorylation of S67. Alternatively, it has also been suggested that S109 phosphorylation can modulate PP2A-B55 inhibitory activity of the phospho-S67 form. As yet, these two hypotheses have been formally discarded ^18,19^.

In this study we identify the major molecular determinants controlling the PP2A-B55 inhibitory activity of Arpp19. Moreover, we elucidate the mechanisms by which phospho-S109 controls S67 Arpp19 phosphorylation and thus, PP2A-B55 inhibitory activity. Finally, we have discovered a double feed-back loop between these two phospho-sites that would be required to coordinate the proper temporal pattern of Arpp19-dependent PP2A-B55 inhibition and Cyclin B/Cdk1 activation, hence ensuring a correct progression through meiosis and mitosis.

## RESULTS

### Aromatic and acidic residues flanking the Gwl phosphorylation site of Arpp19 are essential for a slow dephosphorylation of this site and for the tight binding of this protein to PP2A-B55

Arpp19 is an intrinsically disordered protein that potently inhibits PP2A-B55. Despite its major role in the control of cell cycle progression ^3–6,21^, little is known about the mechanisms by which this protein modulates phosphatase activity. Our laboratory and others have demonstrated that the binding and the inhibition of PP2A-B55 by Arpp19 requires its phosphorylation at S67 by Gwl. This phosphorylation site is located in a motif that is highly conserved from yeast to humans (QKYFDSGDY), which we will refer to from now as the DSG motif (from now) ^3,16,22,23^. Once phosphorylated, Arpp19 acts as an “unfair” substrate that tightly binds PP2A-B55 but that is slowly dephosphorylated by this phosphatase ^16^. Why S67 dephosphorylation of Arpp19 is so slow is unknown. We sought to determine the properties contributing to the inhibitory activity of Arpp19. To this end, we used the well-established cell-free extract system of Xenopus oocytes to perform a structure/function study. In this model, we measured the three major properties of this inhibitor: (1) Gwl site dephosphorylation kinetics, (2) binding to PP2A-B55 and (3) physiologic capacity to promote mitotic entry.

Previous data proposes that the dephosphorylation of ENSA, another member of the endosulfine family, is modulated by the presence of basic KR residues flanking the DSG motif. These residues have been proposed to be a PP2A-B55 recognition signal that would ensure binding to PP2A-B55 and timely ENSA dephosphorylation ^24^.

In order to determine whether dephosphorylation and binding of Arpp19 to PP2A-B55 are modulated by basic KR residues we mutated into alanine three of the basic aminoacids flanking the DSG motif of Arpp19 isoform 1 cloned from Stade VI oocytes. This isoform displays four additional aminoacids at position 62-65 and consequently, residue numbering will differ accordingly in the present study. We constructed a triple alanine mutant of the K36/K38/R40 region (named hereafter as the KKR motif) which was further phosphorylated “*in vitro*” by a recombinant Gwl (Figure 1A). Phosphorylated wildtype and triple KRR mutant Arpp19 were then supplemented to interphase Xenopus egg extracts without ATP to maintain kinases inactive (hereafter called kinase-inactivated extracts). The dephosphorylation pattern of the Gwl site (S71 of Arpp19 isoform 1) was then followed by autoradiography to directly measure phosphatase activity (Figure 1B). As previously reported for ENSA in human cells ^24^, S71 dephosphorylation of wildtype Arpp19 started at 6 minutes upon its addition to kinase-inactivated extracts and fully disappeared at 20 minutes. This dephosphorylation is catalyzed by PP2A-B55 since it was fully blocked upon B55 depletion (Figure 1B). Unlike wildtype Arpp19, S71 dephosphorylation of the KKR triple mutant was not observed throughout the experiment indicating that these basic residues positively modulate S71 dephosphorylation. We next checked whether, as previously proposed, the loss of basic residues could decrease Arpp19 dephosphorylation rate by interfering with PP2A-B55 interaction ^24^. Unexpectedly, PP2A-B55 association to KKR triple mutant was significantly increased compared to wildtype Arpp19 (Figure 1C). Finally, as expected from the decreased S71 dephosphorylation, this mutant kept a full PP2A-B55 inhibitory capacity and promoted mitotic entry when added to Arpp19-depleted egg extracts as indicated by the phosphorylation of Gwl and the loss of the inhibitory phosphorylation of Cdk1 tyrosine 15 (Figure 1D). This data indicates that diminished S71 dephosphorylation in the KKR mutant is not the result of decreased binding to PP2A-B55, and suggests that the KKR motif could directly stimulate S71 dephosphorylation.

**Figure 1.**
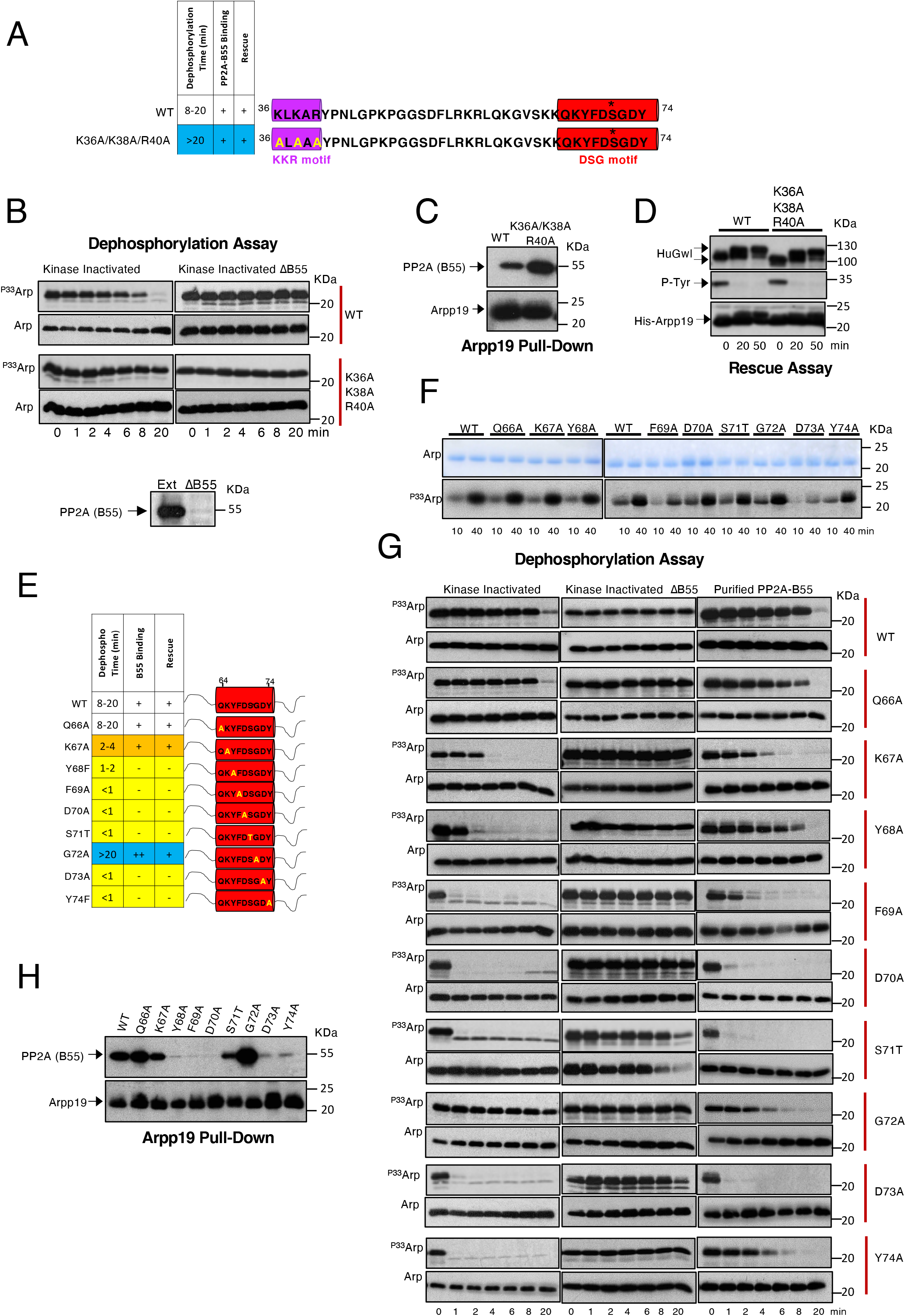
Aromatic and acidic residues flanking S71 Gwl site are essential for slow Arpp19 dephosphorylation rate and tight binding to PP2A-B55. **(A)** Xenopus Arpp19 protein sequence position 36 to 74 including the DSG (red) and the KKR (violet) motifs and the different alanine mutants. Table depicting in the wildtype and the K36A/K38A/R40A Arpp19 mutant the S71 dephosphorylation time, B55-Arpp19 interaction and the capacity to promote mitotic entry in Arpp19-depleted extracts. Blue-colored line highlights S71 Arpp19 dephosphorylation time greater than 20 min. **(B)** 50 ng of wildtype or K36A/K38A/R40A triple Arpp19 mutant phosphorylated “*in vitro*” by GwlK72M were supplemented to kinase-inactivated extracts depleted or not of the B55 protein. The levels of Arpp19 and S71 phosphorylation were analyzed by western blot and autoradiography respectively. B55 levels in kinase-inactivated Xenopus egg extracts after B55 depletion is also shown. **(C)** 20 ng of wildtype or K36A/K38A/R40A triple mutant of a His-Arpp19 pulldown was submitted to western blot and the amount of B55 and the levels of Arpp19 bound to the beads shown. **(D)** Arpp19-depleted extracts were supplemented with human GwlK72M and a wildtype or a triple K36A/K38A/R40A Arpp19 mutant and Human Gwl phosphorylation, Tyr15 of Cdk1 and Arpp19 ectopic levels (His-Arpp19) was assessed. **(E)** A schematic of DSG regions indicating residues mutated into alanine or threonine. Table representing results on the S71 dephosphorylation time, and the capacities to bind B55 or to restore mitotic entry in Arpp19-depleted egg extracts. Yellow, orange and blue lines represent Arpp19 mutants displaying a S71 dephosphorylation time of less than 1 to 2 minutes, between 2 to 4 minutes or more than 20 minutes respectively. **(F)** Wildtype Arpp19 and the indicated mutants of the DSG motif “*in vitro*” phosphorylated by hGwlK72M and 1 μ-sample removed at the indicated time-points to measure S71 phosphorylation by autoradiography (^P33^Arp). The amount of Arpp19 was assessed by Coomassie blue staining. **(G)** S71 dephosphorylation of wildtype Arpp19 or of the indicated DSG Arpp19 mutants was measured in kinase-inactivated extracts depleted or not of B55 as well as upon the addition of a purified PP2A-B55 phosphatase by autoradiography (^P33^Arp). The amount of Arpp19 in each sample was assessed by western blot (Arp). **(H)** B55 levels associated to 20 ng of wildtype or the indicated DSG mutants of His-Arpp19-pulldowns. Arpp19 level in these pull-downs are also shown. All data shown are representative of three different experiments.

We subsequently investigated the impact of mutating Arpp19 in the DSG motif, a sequence that is fully conserved in the endosulfine family of proteins among all the species. Each aminoacid of this motif was mutated into alanine except for the Gwl site S71 whose alanine mutation results in the complete loss of PP2A-B55 binding ^3^. Instead, we substituted this site by threonine, an aminoacid to which PP2A-B55 displays an inherent preference ^25^ (Figure 1E). We could not find differences in the “*in vitro*” capacity of Gwl to phosphorylate S71 in most of the mutants except for F69A and D73A whose phosphorylation was fairly decreased (Figure 1F). This data disagrees with a previous report suggesting a dramatic drop in the phosphorylation of this Arpp19 site by Gwl when aromatic and acidic residues as well as S71 itself were mutated in human cells ^24^. We do not know the reason for these differences. However, it is possible that the decline of S71 phosphorylation observed in these Arpp19 mutants could result from increased PP2A-B55-dependent dephosphorylation instead of decreased Gwl-dependent phosphorylation.

To test this hypothesis, we thus measured S71 dephosphorylation of “*in vitro*” phosphorylated mutants in kinase-inactivated egg extracts. The level of recombinant Arpp19 protein was evaluated by western blot (Arp) whereas the rate of S71 dephosphorylation was captured by autoradiography (^P33^Arp). Results are shown in Figure 1G and represented in the table of Figure 1E. Interestingly, except for Q66A and G72A that kept normal or decreased dephosphorylation rates respectively, all the other mutants of Arpp19 were rapidly (for K67A) or a very rapidly (for the rest) dephosphorylated on S71. In agreement with Figure 1B, this dephosphorylation pattern is promoted by PP2A-B55 since it was fully blocked by B55 depletion in these extracts and is phenocopied in most of the mutants when “*in vitro*” dephosphorylation assays were performed using a purified PP2A-B55 (Figure 1G). Additionally, the S71 dephosphorylation rate correlated with the ability of these mutants to bind B55 with normal or an increased association for mutants displaying a regular or a slower dephosphorylation rate (Q66A and G72A respectively), intermediate association for mutants with a moderate dephosphorylation (K67A), and with fully loss of B55 binding for mutants with a far faster dephosphorylation rate (Y68A, F69A, D70A, D73A, Y74A) (Figure 1H). S71T Arpp19 represents an exception to this rule, since it displayed a fast dephosphorylation but a partial binding to B55. This result is not surprising as the catalytic efficiency of PP2A-B55 is known to be 20-fold higher for phosphorylated threonine than serine ^25^. Interestingly, the most rapid dephosphorylation was observed when the acidic and aromatic residues of the DSG motif were mutated, the same mutants previously proposed to lose Gwl-dependent phosphorylation in human cells. Hence, our results confirm that the weak phosphorylation of these mutants at S71 results from their dephosphorylation by PP2A-B55 and not from a decreased phosphorylation by Gwl ^24^.

As expected, rapid dephosphorylation of S71 and decreased binding to B55 in these mutants was also associated with their incapacity to promote mitotic entry upon their addition to Arpp19-depleted egg extracts (Supplementary Figure 1B) indicating that although they still behave as PP2A-B55 substrates, they lost their phosphatase inhibitory activity.

We then sought to assess how these mutations could promote rapid dephosphorylation and loss of PP2A-B55 binding. One hypothesis is that these mutations could exclusively impact the catalysis of S71 promoting a faster dephosphorylation of S71. As a result, the residence time of Arpp19 on PP2A-B55 enzyme would be reduced, a process that would explain the loss of this interaction in our pulldown analysis. A second hypothesis would involve not only of S71 dephosphorylation reaction but also changes in the local conformation of Arpp19 that would modify Arpp19-PP2A-B55 interaction. In order to discriminate between these two possibilities, we blocked S71 catalysis by “*in vitro*” thio-phosphorylation and further monitored the inhibitory activity of these mutants toward PP2A-B55 (Supplementary Figure 1C). When constitutively phosphorylated at S71 none of these mutants were able to promote mitotic entry, suggesting that, aside its putative effect on the catalysis of the Gwl site, these mutations might affect Arpp19-PP2A-B55 interaction.

### The C-terminus of Arpp19 controls PP2A-B55 inhibition by modulating S71 dephosphorylation rate

To ask whether sequences others than the DSG motif could be involved in the PP2A-B55 inhibitory activity of Arpp19, we mutated the N- and the C-terminus of this protein (Figure 2A). N-terminal deletion mutant did not modify either S71 dephosphorylation (Figure 2B) or its inhibitory capacity (Supplementary Figure 2A) and only slightly decreased PP2A-B55 association (Figure 2C). Conversely, the (1-75) Arpp19 mutant lacking most of the C-terminal region displayed a very fast dephosphorylation of the S71 by PP2A-B55 (less than 1 minute) (Figure 2D, top panel). This dephosphorylation pattern was associated with a loss of its interaction with PP2A-B55 (Figure 2E) and with its incapacity to promote mitotic progression in Arpp19-depleted egg extracts (Supplementary Figure 2B). We thus constructed three shorter C-terminal mutants (1-86; 1-93; 1-101) (Figure 2A). All these mutants also displayed an acceleration of PP2A-B55-dependent S71 dephosphorylation (Figure 2D) as well as a loss of the interaction with this phosphatase (Figure 2E). Accordingly, they were unable to promote mitotic entry although this capacity was rescued when S71 was thio-phosphorylated (Supplementary Figure 2B and 2C respectively). Since the ability of these mutants to inhibit PP2A-B55 is lost but can be restored by the stable phosphorylation of S71, this data suggests that the C-terminus of Arpp19 modulates its phosphatase inhibitory capacity by exclusively modifying the ability of PP2A-B55 to catalyze S71 dephosphorylation, with no major impact on Arpp19-PP2A-B55 association.

**Figure 2.**
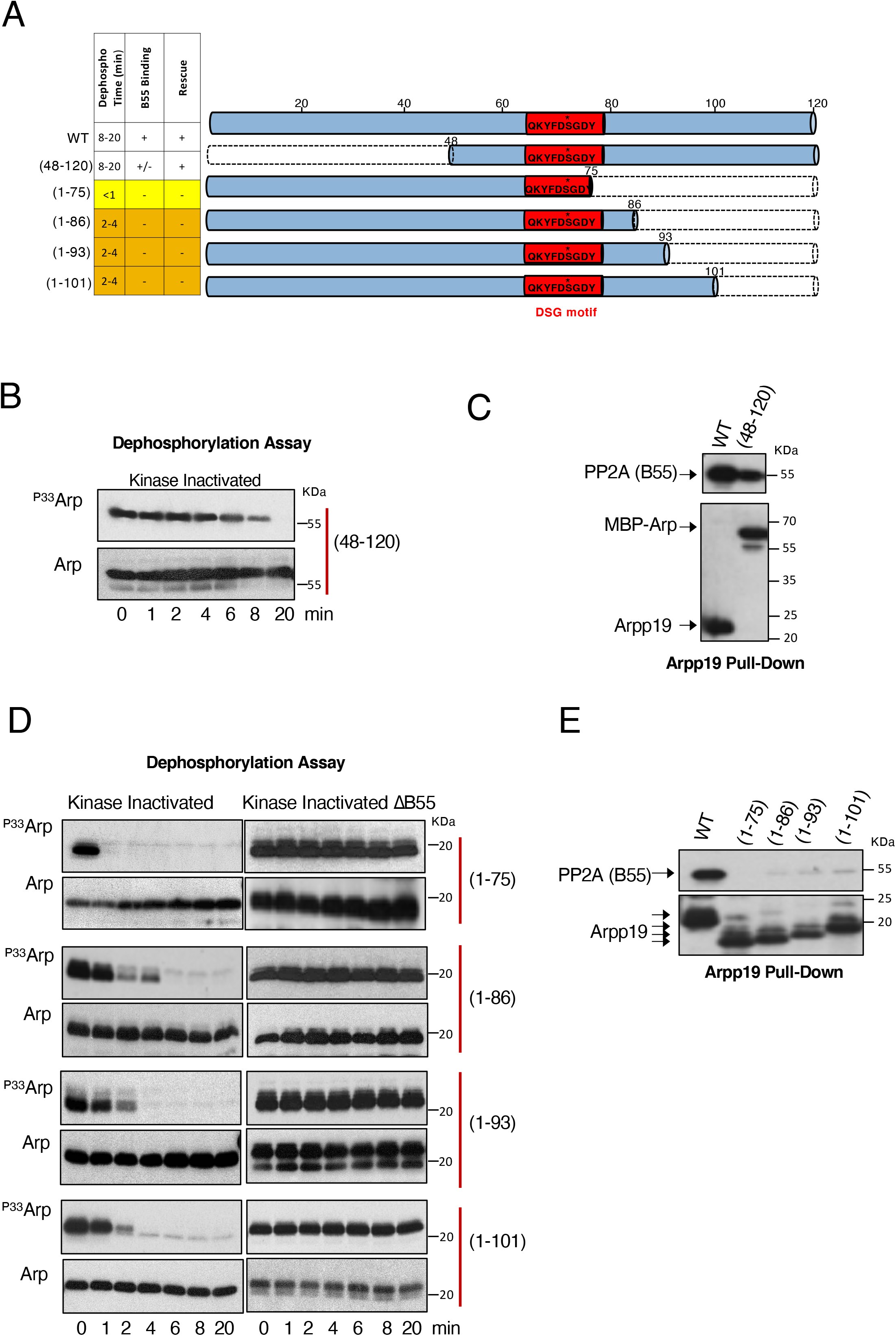
The C-terminus of Arpp19 controls PP2A-B55 inhibition by modulating S71 dephosphorylation rate. **(A)** Represented in dashed lines are the sequences deleted in the specified Arpp19 mutants. A table summarizing data of the S71 dephosphorylation time in kinase-inactivated extracts as well as the binding or not to B55 and the capacity to restore the mitotic state in Arpp19-depleted extracts of all these mutants is also shown. Yellow and orange lines denote a dephosphorylation time of S71 of less of 1 minute or between 2 to 4 minutes respectively. **(B)** The dephosphorylation of S71 of (48-120) Arpp19 mutant was assayed in kinase-inactivated extracts and revealed by autoradiography. The amount of this Arpp19 mutant form in each sample is also shown. **(C)** A volume of His-Arpp19 pulldown sample corresponding to 20 ng of wildtype or (48-120) mutant is submitted to western blot and the associated B55 protein as well as the amount of Arpp19 present in the beads shown. Due to the insolubility of the His-Arpp19 (48-120) mutant, we used a double tagged MBP-His Arpp19. **(D)** S71 dephosphorylation assay of the indicated Arpp19 mutants was tested in kinase-inactivated extracts depleted or not of B55 and revealed by autoradiography. The amounts of Arpp19 mutant proteins present in each sample was checked by western blot. **(E)** Levels of B55 associated to a volume of beads equivalent to 20 ng of the wildtype and the indicated C-terminal Arpp19 mutants. The amount of these mutants bound to the beads is also shown. All data are representative of three different experiments.

We next constructed shorter C-terminal mutants in which regions 78 to 95 or 96 to 111 were deleted (Figure 3A). Deletion of each of these sequences promoted acceleration of PP2A-B55-dependent S71 dephosphorylation (Figure 3B), loss of the interaction with this phosphatase (Figure 3C) and the inability to promote mitotic entry (Figure 3A and Supplementary Figure 2D). We next checked the conservation of these two sequences between endosulfines and among species. This analysis revealed a motif highly conserved at positions from 96 to 111 (Supplementary Figure 2E), a motif that we named as the “cassette motif” (Figure 3A). We thus constructed two different mutants in which two (T103A/P104A; 2A mutant) or six conserved aminoacids (T103A/P104A/D106A/L107A/P108A/Q109A; 6A mutant) were mutated into alanine. Interestingly, the 2A mutant (Figure 3B, “2A”) displayed an accelerated S71 dephosphorylation that was more pronounced in the 6A mutant (Figure 3B, “6A”). Moreover, PP2A-B55 binding was decreased for 2A and lost for 6A mutant (Figure 3C). Finally, the 2A but not 6A mutant conserved its ability to trigger mitosis (Supplementary Figure 2F). However, this faculty was rescued in the latter mutant when S71 was thio-phosphorylated (Supplementary Figure 2G), hence confirming that the C-terminal cassette motif of Arpp19 is essential to confer correct dephosphorylation catalysis of the Gwl site. Moreover, it demonstrates that this capacity is sequence specific.

**Figure 3.**
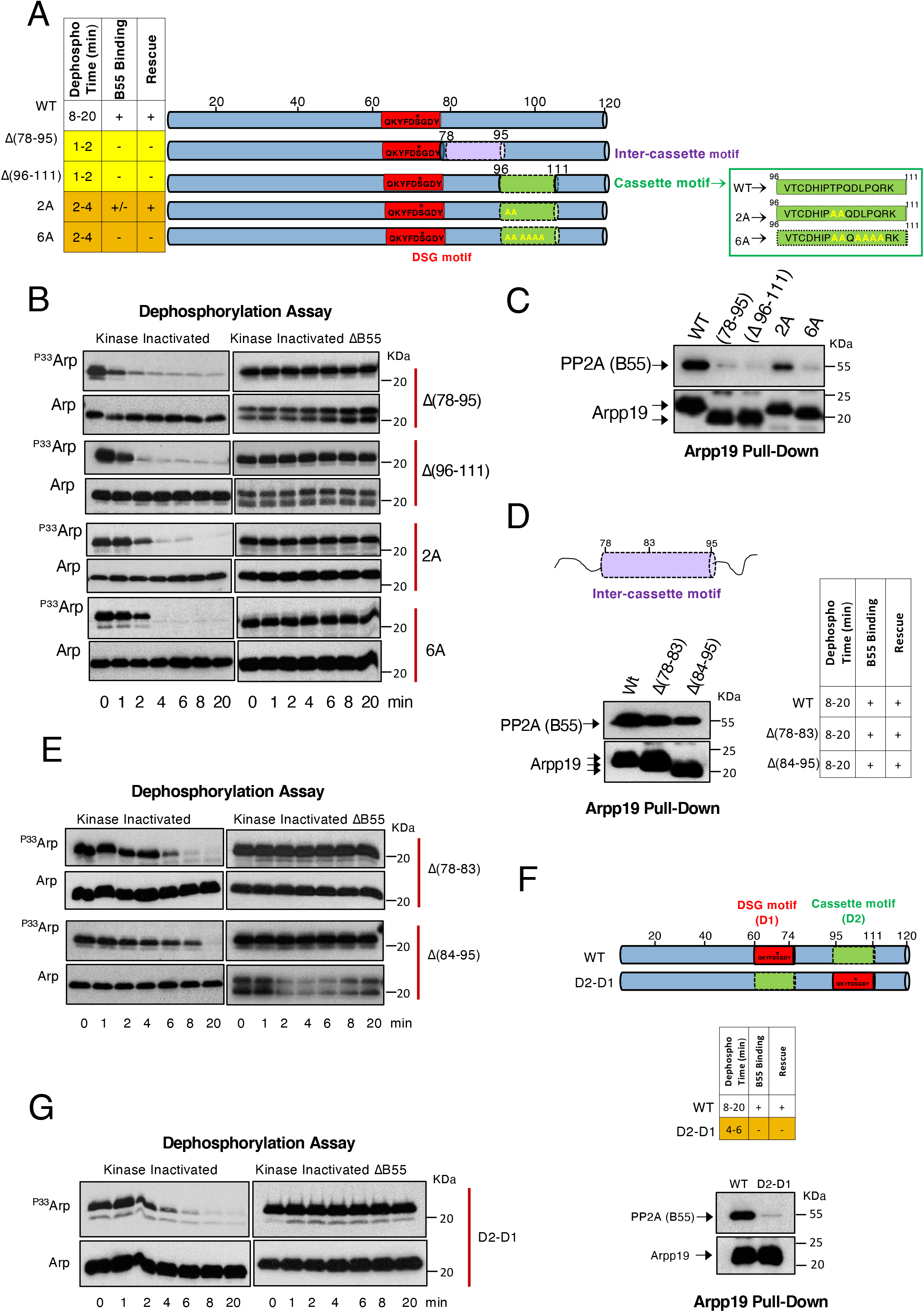
A specific sequence on the cassette motif of Arpp19 and a critical distance between this region and the DSG motif are essential for the correct timing of S71 dephosphorylation by PP2A-B55 **(A)** Schematic representation of the “inter-cassette” (sequence 78 to 95) and the “cassette” (sequence 96 to 111) deleted regions of Arpp19 as well as the residues on the “cassette motif” that have been mutated into alanine. A table summarizing data on the timing of S71 dephosphorylation in kinase-inactivated extracts, the association or not of these mutants to B55 and their capacity to restore mitosis in Arpp19-devoid extracts is shown. **(B)** Dephosphorylation rate of S71 of the indicated mutants of Arpp19 in kinase-inactivated extracts devoid or not of B55. The amount of Arpp19 in each sample was assessed by western blot. **(C)** Western blot showing the association of B55 to the indicated mutants of Arpp19. **(D)** Scheme depicting the two regions that have been deleted in the “inter-cassette” motif. Table illustrating data on S71 dephosphorylation timing as well as the capacity to bind B55 and to restore mitosis of the two mutants. Western blot showing the amount of B55 present in the His-App19 pull down assays of the indicated Arpp19 mutant forms. **(E)** S71 dephosphorylation assays of the indicated mutants of Arpp19 in kinase-inactivated extracts devoid or not of B55. The amount of Arpp19 in each sample was assessed by western blot. **(F)** A schematic of the DSG (D1) and the cassette (D2) regions of Arpp19 that have been exchanged in the D2-D1 mutant. A table with the dephosphorylation, binding and rescue results is also shown. The association of B55 to the D2-D1 mutant compared to the wildtype Arpp19 is shown. **(G)** Dephosphorylation assay of S71 of the wildtype and the D2-D1 mutant in kinase-inactivated extracts that have been depleted or not of B55. Representative data of at least three different experiments.

Unlike the cassette motif, the sequence from 78 to 95, (called hereafter as the “inter-cassette motif”) (Figure 3A and D) appears to be less conserved (Supplementary Figure 2E), yet, we decided to test whether the length instead of the sequence identity of this region would be important for Arpp19 to preserve its PP2A-B55 binding and inhibitory activity. We thus constructed two mutants in which either region 78 -83 or region 84-95 were deleted (Figure 3D, pull-down). Both of these deletions promoted a normal dephosphorylation of S71 (Figure 3E), a standard interaction with PP2A-B55 (Figure 3D) and mitotic entry in Arpp19-depleted egg extracts (Supplementary Figure 2H). This indicates that no specific sequence in these zones is required but instead a minimal length of the inter-cassette motif is critical. Thus, a distance between the DSG and the cassette motifs determines the correct inhibitory activity of Arpp19.

From this data, we conclude that the presence of two sequence specific regions are essential for Arpp19 inhibitory activity, the DSG (D1) and the cassette (D2) motifs (Figure 3F). Since Arpp19 is an intrinsically disordered protein, we next wondered whether, beside sequence specificity and lenght, the localization of these two regions in the protein was also important for their physiological function. We thus constructed a chimeric Arpp19 protein in which these two regions were exchanged (Figure 3F). Interestingly, although this chimeric protein was normally phosphorylated in S71, this residue was fastly dephosphorylated (Figure 3G), and was fully unable to bind PP2A-B55 (Figure 3F) and to promote mitosis (Supplementary Figure 2I) even upon thio-phosphorylation of S71 (Supplementary Figure 2J). Hence, the location of these two regions into the protein is critical to maintain the PP2A-B55 inhibitory activity of Arpp19.

### Phosphorylation of S113 modifies secondary structure propensity of the cassette motif and the temporal pattern of S71 dephosphorylation

Our results indicate that the cassette motif is essential for Arpp19 physiological activity by modulating the catalysis of PP2A-B55-dependent dephosphorylation of S71. Interestingly, S113, close to this motif, is phosphorylated by PKA in prophase Xenopus oocytes and this phosphorylation is essential to block meiotic resumption ^18^. During meiotic resumption upon progesterone addition, S113 is partly dephosphorylated. This leads to the activation of Cyclin B/Cdk1, the subsequent phosphorylation of Arpp19 at S71 and Germinal Vesicle Breakdown (GVBD). Due to the close proximity, we hypothesized that S113 phosphorylation could modulate the cassette motif and thus, have an impact on S71 dephosphorylation and PP2A-B55 inhibitory activity. To investigate this hypothesis, we performed a [^1^H-^15^N] HSQC Nuclear Magnetic Resonance (NMR) spectrum of non-phosphorylated Arpp19 and a S113D phospho-mimetic mutant. Both spectra confirmed an unfolded state of this protein (Figure 4A). When superimposed, most of the peak positions of the spectra remained essentially unchanged. However, nine residues within the adjacent regions to S113 displayed a variation of chemical shift. From those residues, three were a part of the cassette motif (positions 109, 110 and 111) and six were located after S113 (positions from 114 to 119). This suggests that S113 phosphorylation could modify the temporal pattern of PP2A-B55-dependent dephosphorylation of S71 by impacting the secondary structure of the cassette motif.

**Figure 4.**
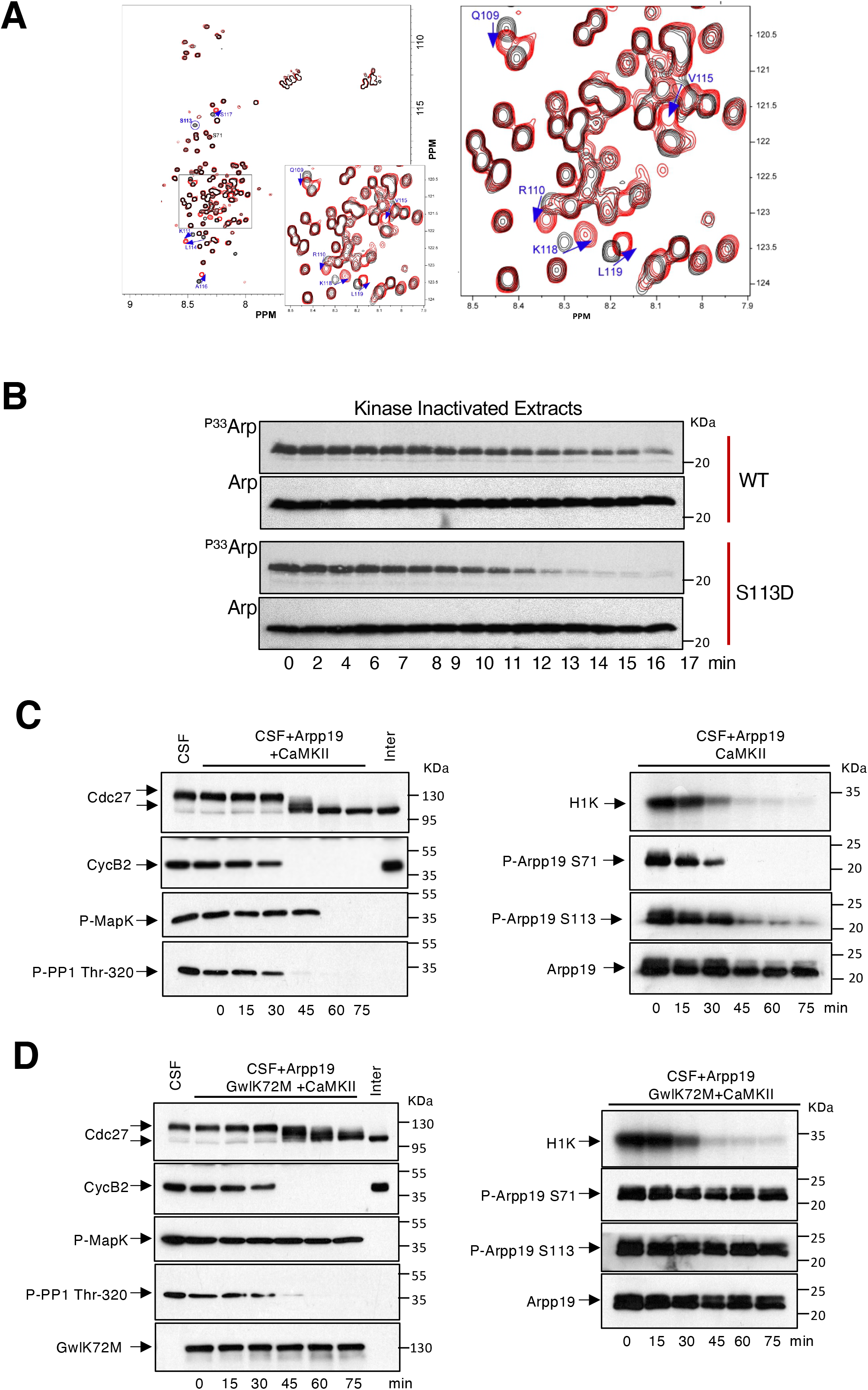
Phosphorylation of S113 modifies secondary structure propensity of residues on the “cassette” motif and the temporal pattern of S71 dephosphorylation. **(A)** Superposed Arpp19 nuclear magnetic resonance (NMR) fingerprint of wildtype (black) and S113D (red) Arpp19 mutant. [^1^H-^15^N] HSQC spectrum of Arpp19 from Xenopus oocytes at 700 MHz, pH 6.8, recorded at 20°C on a 1 mM ^15^N-uniformly labeled sample dissolved in 10 mM Na-Phosphate buffer with 100 mM NaCl. Cross-peak assignments are indicated using the one-letter amino acid and number. The central part of the spectrum is expanded in the insert and shown in the right panel. Blue arrows denote chemical shifts of the indicated residues. **(B)** Wildtype and S113D Arpp19 mutant were phosphorylated “*in vitro*” with of [γ^33^P] ATP by GwlK72M on S71 and supplemented to kinase-inactivated Xenopus egg extracts. The dephosphorylation of this residue was analyzed at the indicated times by autoradiography (^P33^Arp) and the amount of Arpp19 in each sample measured by western blot (Arp). **(C)** CSF egg extracts were supplemented with a trace level of Arpp19 purified protein and activated to exit meiosis by the addition of active CamKII. The levels and dephosphorylation of the indicated proteins were analyzed by western blot whereas Cyclin B/Cdk1 activity was measured by histone H1 phosphorylation (H1K). “Inter” denotes interphase egg extracts. **(D)** As for (C) except that S71 Arpp19 dephosphorylation and PP2A-B55 reactivation upon meiosis exit in these extracts was blocked by the concomitant addition of GwlK72M purified protein. Data shown in the figure are representative of three different experiments.

Accordingly, S71 dephosphorylation of Arpp19 was significantly faster in S113D Arpp19 mutant compared to wildtype Arpp19 in kinase-inactivated extracts (Figure 4B).

Altogether, these results establish that S113 phosphorylation is a key site that modulates PP2A-B55 inhibitory activity of Arpp19.

While, PKA has been characterized as the kinase promoting the phosphorylation of this residue ^17,26,27^, nothing is known about the counterbalancing phosphatase. We thus, focused on identifying this phosphatase.

To this end, we checked S113 and S71 dephosphorylation using a trace amount of Arpp19 in CSF extracts. These extracts were then forced to exit meiosis by the addition of active CamKII. Both S71 and S113 were fully dephosphorylated at meiotic exit, although S71 dephosphorylation was significantly faster than S113 (Figure 4C). Interestingly, when S71 phosphorylation was maintained by adding a Gwl hyperactive form, cyclin B degradation and CSF exit was normally performed (Figure 4D, left panel). However, S113 dephosphorylation was not observed strongly suggesting that this dephosphorylation depends on PP2A-B55 activity (Figure 4D, right panel).

### Dependence of S113 dephosphorylation of Arpp19 on the phosphorylation of the S71

To further elucidate whether PP2A-BB5 is responsible of S113 dephosphorylation, we used kinase-inactivated extracts supplemented with either a phospho-S113 or a phospho-S71 Arpp19 protein. As expected from the absence of kinase activity in these extracts, dephosphorylation of both S113 and S71 of Arpp19 was drastically accelerated (2 to 6 minutes) (Figure 5A) when compared to the one observed in CamKII-treated CSF extracts (30 to 45 minutes) (Fig 4C). Moreover, S113 was dephosphorylated before S71 (Figure 5A). Interestingly, this inversed order of dephosphorylation was no longer observed when extracts were simultaneously supplemented with both S71 and S113 phosphorylated Arpp19 forms (Fig 5B). These results indicate that S71 site modulates S113 dephosphorylation.

**Figure 5.**
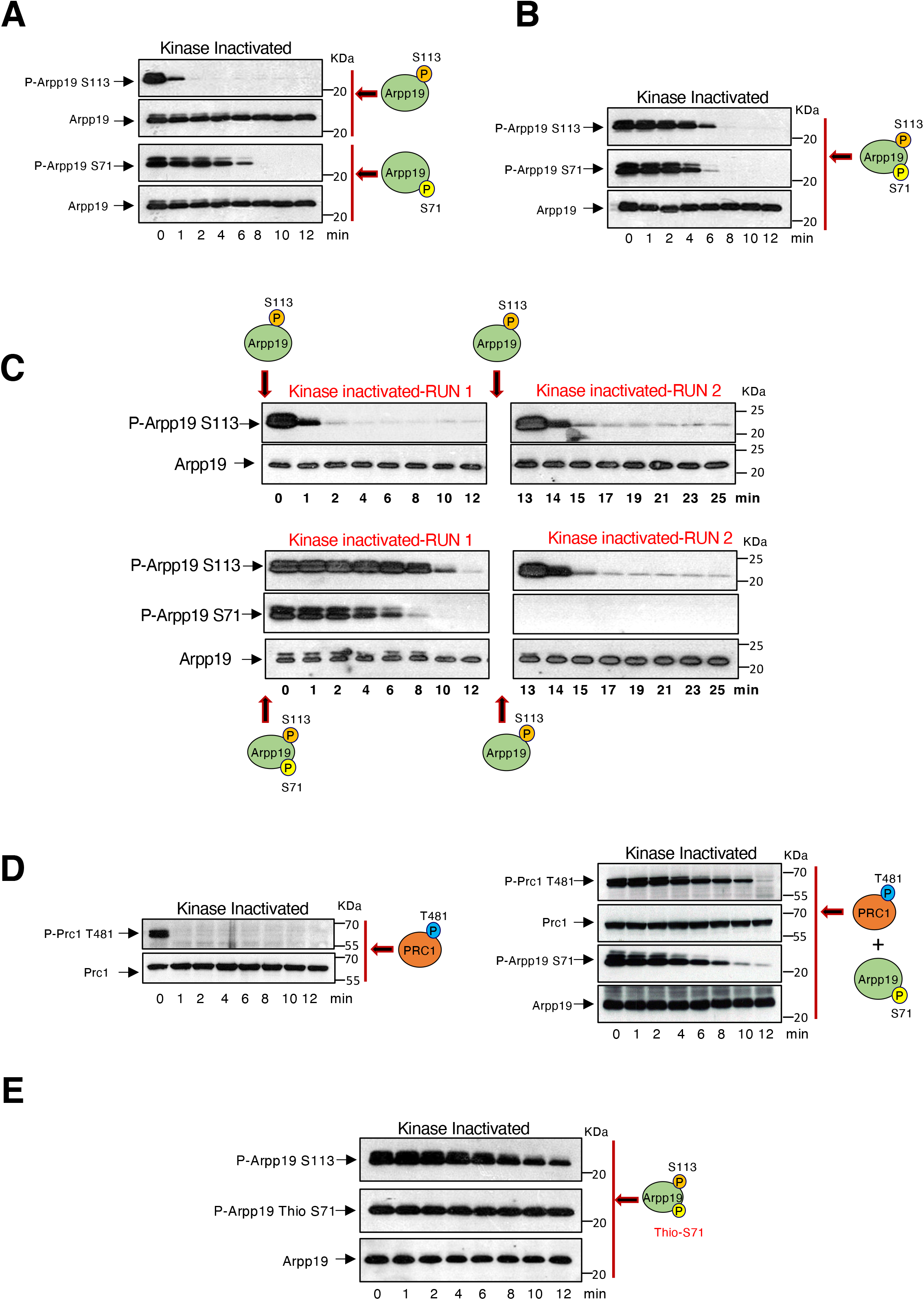
Dependence of S113 dephosphorylation on the phosphorylation of S71 of Arpp19. **(A)** Arpp19 phosphorylated “*in vitro*” in S71 or S113 residues by Gwl or PKA respectively, were separately supplemented to kinase-inactivated Xenopus egg extracts and the dephosphorylation rate of each site as well as the total amount of this protein were analysed by western blot. **(B)** Arpp19 “*in vitro*” phosphorylated by Gwl and Arpp19 “*in vitro*” phosphorylated PKA were mixed together into kinase-inactivated extracts and the dephosphorylation of each of these sites analyzed at the indicated time-points. **(C)** In a first run of dephosphorylation, a pulse of Arpp19 phosphorylated by PKA or by both PKA and Gwl was supplemented to kinase-inactivated extracts. Samples were then recovered at the indicated time-points. After 12 minutes, a second round of dephosphorylation was performed in these extracts upon the re-addition of a new pulse of Arpp19 phosphorylated “*in vitro*” by PKA on S113. Phosphorylation of S71 and S113 in the samples were evaluated by western blot with specific phospho-antibodies. **(D)** Prc1 was phosphorylated “*in vitro*” by purified Cyclin A/Cdk and supplemented alone (left panels) or together with phospho-S71 Arpp19 (right panels) to kinase-inactivated extracts and the phosphorylation of T481 of Prc1 and S71 of Arpp19 as well as the amount of these two proteins were examined by western blot. **(E)** Arpp19 phosphorylated “*in vitro*” by PKA on S113 and thio-phosphorylated on S71 by GwlK72M were mixed into kinase-inactivated extracts. Dephosphorylation of these two residues were then measured at the indicated time-points by western blot. Data shown in the figure is representative of at least three different experiments.

To deeply investigate this hypothesis, we used two kinase-inactivated extract samples. In one of this sample, phospho-S113 Arpp19 was added, whereas the other one was supplemented with both phospho-S71 and phospho-S113 Arpp19 forms. Dephosphorylation of S113 and S71 was then followed during 12 min (Figure 5, RUN 1). After this period of time, both extract samples were supplemented again with phospho-S113 Arpp19 and the dephosphorylation of this residue was recorded for an additional period of 12 minutes (Figure 5, RUN 2). Interestingly, dephosphorylation of S113 was very fast (about 1 to 2 minutes) in both runs in the extract sample where only phospho-S113 Arpp19 was supplemented. Conversely, during the first run, a dependence of dephosphorylation of S113 on S71 dephosphorylation was observed in the extract sample in which phospho-S71 and phospo-S113 were simultaneously added. However, in this sample S113 dephosphorylation acquired again a fast dephosphorylation rate during the second run when phospho-S71 Arpp19 was absent. This data confirms a regulatory role of S71 phosphorylation on S113 dephosphorylation.

Phospho-S71 Arpp19 is known to be an “unfair*”* substrate of PP2A-B55 that negatively regulates this enzyme by competition ^16^. It is thus possible that as for S71, dephosphorylation of S113 could be catalyzed by PP2A-B55 and consequently, be slowed down when Arpp19 is phosphorylated at S71. To test this hypothesis, we first compared the dephosphorylation pattern of another well-known substrate of PP2A-B55, the residue T481 of Prc1, in the presence or absence of phospho-S71 Arpp19. T481 Prc1 dephosphorylation is very rapid in the absence of phospho-S71 Arpp19 (Fig 5D). However, phospho-T481 was maintained until S71 was fully dephosphorylated when a phospho-S71 Arpp19 form was added and displayed an identical timing of dephosphorylation than S113 when both S71 and S113 Arpp19 forms are present in the extract (compare Fig 5C, Run1, low left panel and Figure 5D, right panel). Finally, we assessed the impact on S113 dephosphorylation of a S71-thiophosphorylated Arpp19, known to strongly inhibit PP2A-B55. As expected, S113 phosphorylation was mostly stable under these conditions (Fig 5E), indicating that Arpp19 dephosphorylation at S113 mostly relies on PP2A-B55.

### PP2A-B55 is the phosphatase responsible for the dephosphorylation of Arpp19 S113

The data reported above strongly suggest a role of PP2A-B55 in the dephosphorylation of S113 of Arpp19. To fully elucidate the identity of this phosphatase we performed a biochemical approach in which kinase-inactivated extracts were fractionated by gel chromatography using a Superdex 200 column. The different fractions were then tested for S113 Arpp19 dephosphorylation activity using recombinant Arpp19 phosphorylated *“in vitro”* on either S113 or on S71 as a substrate. S113 and S71 dephosphorylation activities were both recovered at fractions 32-42 overlapping with the peak of PP2A-A, C, B55 and B56 abundance supporting again the hypothesis of PP2A being responsible for these dephosphorylations (Fig 6A). Although both B55 and B56 were present in S113 dephosphorylation active fractions, we decided to focus on the putative involvement of PP2A-B55 given our results obtained in extracts using S71-thio-phosphorylated Arpp19. We first monitored the stability of S113 phospho-site in B55 immunodepleted interphase extracts. As expected, both T481 of Prc1 and S71 of Arpp19, two known substrates of PP2A-B55, were not dephosphorylated in B55 depleted interphase extracts (Figure 6B). Similarly, S113 dephosphorylation was dramatically delayed in kinase-inactivated extracts devoid of B55 regardless of the presence of a S71 phosphorylated Arpp19 form indicating again that PP2A-B55 is the main phosphatase involved in dephosphorylation of S113 of Arpp19.

**Figure 6.**
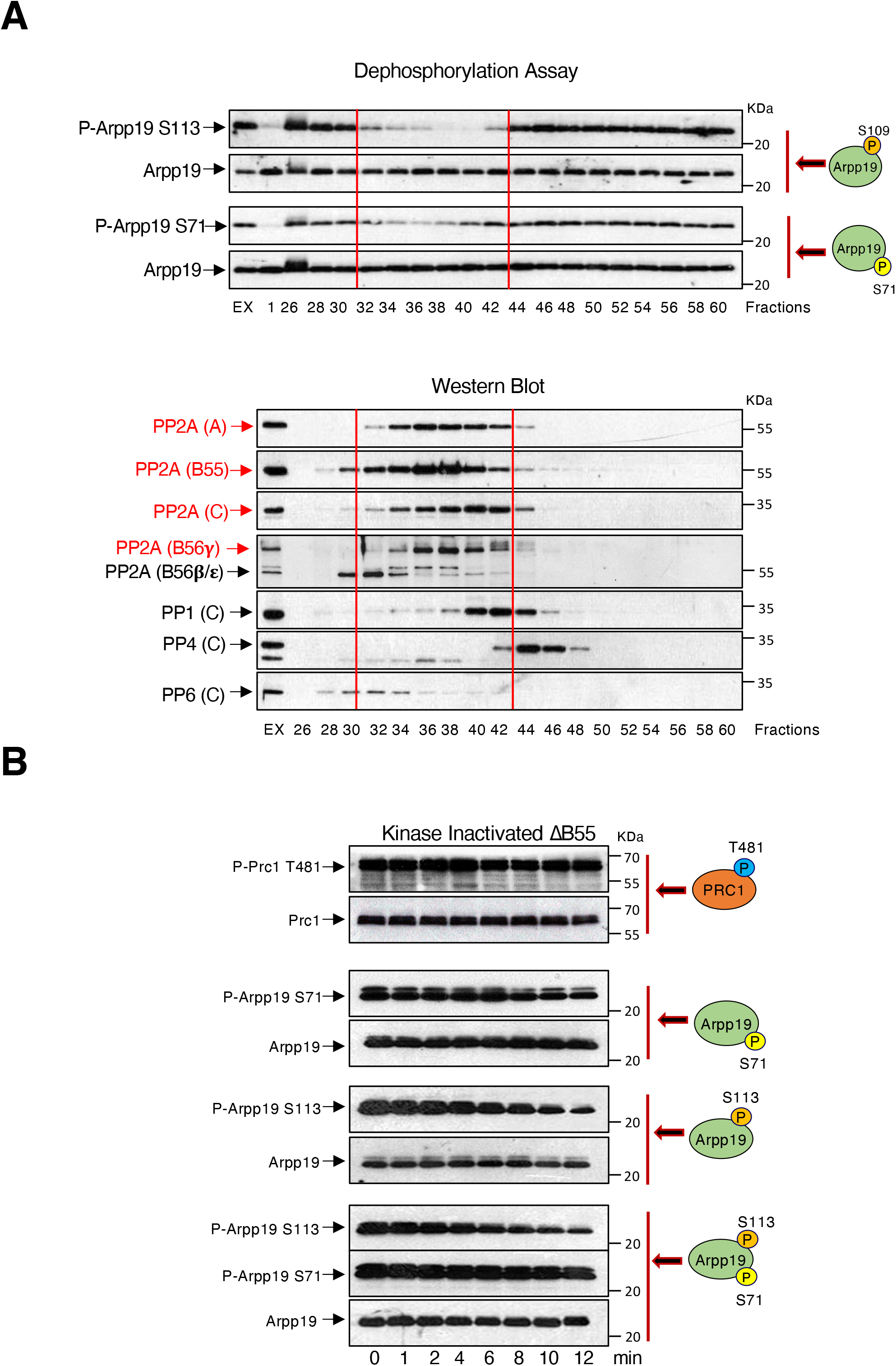
PP2A-B55 is the phosphatase responsible of the dephosphorylation of Arpp19 on S113. **(A)** Kinase-inactivated Xenopus egg extracts were submitted to gel chromatography and eluted fractions subsequently supplemented with Arpp19 phosphorylated “*in vitro*” on either S113 or S71. The dephosphorylation of these residues as well as the amount of Arpp19 were then measured by western blot (upper panels, dephosphorylation assay). “EX” corresponds to phosphorylation at time 0 of the indicated residues upon directly mixed with a kinase-inactivated extract. The presence of the indicated proteins in elution fractions and in the kinase-inactivated extract sample were assessed by western blot (lower panels, western blot). Red lines highlight the fractions displaying S113 and S71 dephosphorylation activity. The name of the proteins whose level picked in these fractions are also depicted in red. **(B)** Prc1 “*in vitro*” phosphorylated on T481 by purified Cyclin A/Cdk as well as Arpp19 phosphorylated on either S71, or on S113 were supplemented together or separately to kinase-inactivated extracts depleted of B55 and the dephosphorylation of the corresponding residues analyzed over the time by western blot. Data of the figure is confirmed in three different experiments.

We subsequently confirmed this data *“in vitro”* using a purified PP2A-B55 phosphatase (Supplementary Fig 1A and Figure 7A). We assessed the dephosphorylation of the S113 and S71 phosphorylated forms of Arpp19 when separately or simultaneously added to the reaction mix. In accord with our data above, the temporal pattern of S71 dephosphorylation was unchanged under all conditions. On the contrary, S113 phosphorylation disappeared very rapidly in the absence of phospho-S71 but did not occurred upon full S71 dephosphorylation when this phospho-residue was present. Finally, we checked the effect of the double addition to the reaction mix of a phospho-S113 and a thio-phospho-S71 Arpp19 forms. Under these conditions, S113 phosphorylation was stabilized throughout the experiment confirming that PP2A-B55 is the main phosphatase involved in S113 dephosphorylation. Interestingly, this regulatory mechanism is shared and interchangeable by the other PP2A-B55 inhibitor ENSA. Accordingly, dephosphorylation of S109 of ENSA in kinase-inactivated extracts was very fast in a single phosphorylated ENSA protein whereas it was maintained until S67 phosphorylation fully disappeared when both S67 and S109 were phosphorylated (Figure 7B). Similarly, when both phospho-S109 ENSA and phospho-S71 Arpp19 were simultaneously supplied to kinase-inactivated extracts, dephosphorylation of phospho-S109 ENSA was only observed when phospho-S71 Arpp19 fully disappeared (Figure 7C). An identical dephosphorylation timing was followed when phospho-S113 Arpp19 and phospho-S67 ENSA were used.

**Figure 7.**
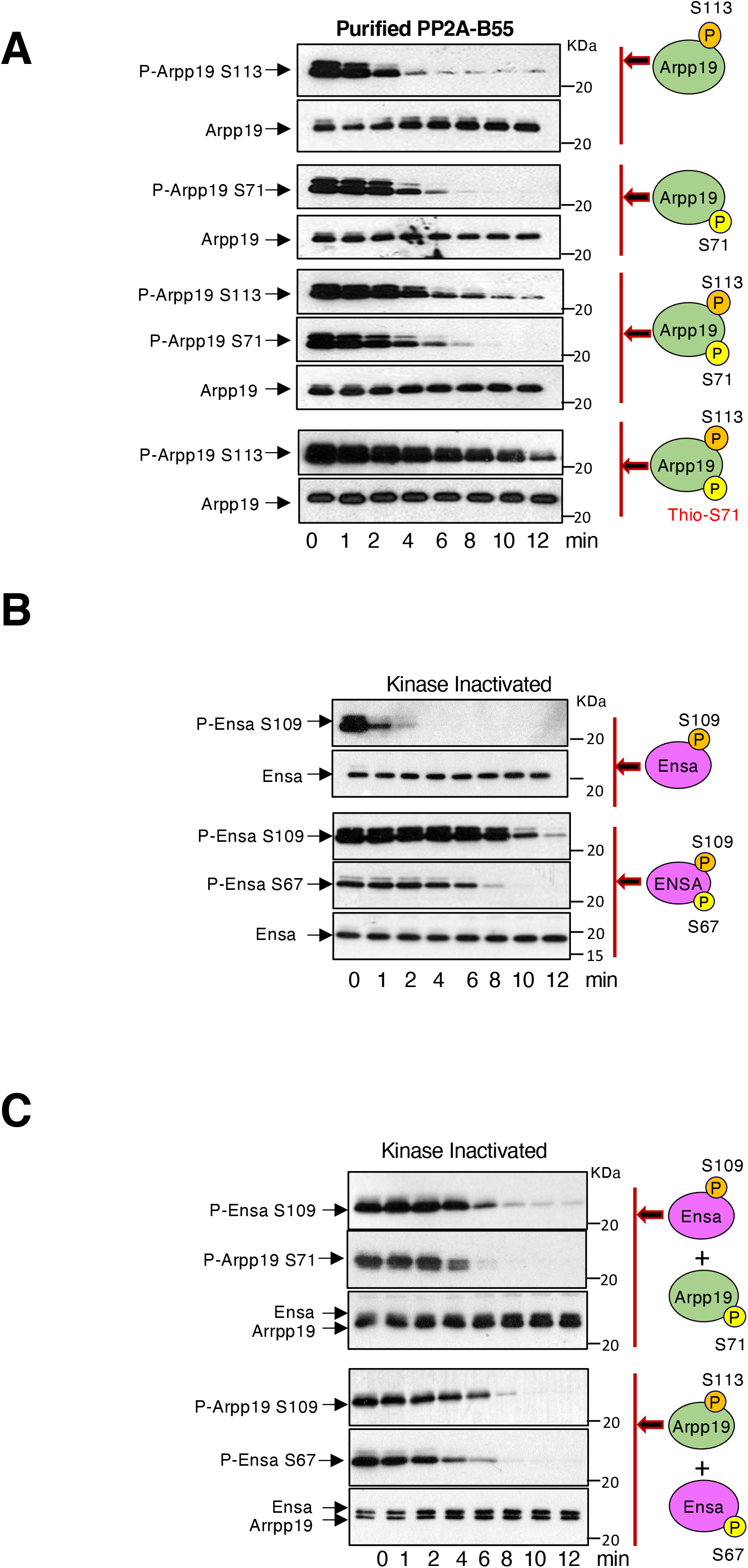
Phospho-S71 negatively regulates dephosphorylation of S113 of Arpp19 by PP2A-B55. **(A)** Arpp19 was phosphorylated “*in vitro*” on S113, S71 or thio-phosphorylated on S71, supplemented alone or with the indicated combinations to kinase-inactivated extracts and the temporal pattern of dephosphorylation analyzed. **(B)** ENSA protein was phosphorylated “*in vitro*” on S67 by GwlK72M or S109 by PKA and supplemented alone or combined as indicated to kinase-inactivated extracts and the dephosphorylation time of these residues as well as the amount of ENSA protein examined. **(C)** Phospho-S109 ENSA and phospho-S71 Arpp19 or phospho-S113 Arpp19 and phospho-S67 ENSA were simultaneously added to kinase-inactivated extracts and the timing of dephosphorylation of the different phospho-sites analyzed. Experiments supporting the data of this figure were performed at least three times.

## DISCUSSION

Arpp19 is a powerful PP2A-B55 inhibitor whose activity is essential for mitotic and meiotic progression. This inhibitory activity of Arpp19 is acquired via its phosphorylation at a unique residue at S67/S71 by Gwl ^3,4^. At mitotic entry, Gwl activation phosphorylates Arpp19. This induces its binding to PP2A-B55 and the inhibition of this phosphatase, hence allowing the stable phosphorylation of Cyclin B/Cdk1 substrates and a normal mitotic progression. Accordingly, the knockout of Arpp19 in MEFs leads to PP2A-B55 reactivation, the precocious dephosphorylation of mitotic substrates and the disruption of the correct temporal order of cellular events during mitotic progression ^6^. Beside its important role in mitosis, the inhibition of PP2A-B55 by Gwl/Arpp19 is also required for Cyclin B/Cdk1 dependent phosphorylation during meiotic division. As so, the microinjection of a S67/S71 phosphorylated Arpp19 protein in prophase I-arrested oocytes triggers meiotic resumption in the absence of progesterone stimulation ^18,21^.

While the essential function of Arpp19-dependent PP2A-B55 inhibition in mitosis and meiosis has been well established, little is known about the molecular mechanisms controlling this inhibition. In this line, it has been demonstrated that ENSA, the other member of the endosulfine family of proteins acts as an “unfair*”* substrate of this phosphatase with a tight-binding to this enzyme and a slow dephosphorylation rate. Thus reducing PP2A-B55 substrate dephosphorylation by competition ^16^. This “unfair” competition model implies a dependence of the inhibitory capacity of these proteins on two main parameters: (1) the ability of Arpp19/ENSA to bind PP2A-B55, reinforced by its Gwl-dependent phosphorylation on S67/S71 and (2) the subsequent capacity of these proteins to dissociate from PP2A-B55 via the dephosphorylation of this residue. It is therefore essential to elucidate the structural properties defining phosphatase interaction and S67/S71 dephosphorylation rate to understand the molecular mechanisms of Arpp19-dependent PP2A-B55 inhibition.

Here, we performed an extended structure/function study to determine these properties. To this end, we set up three different assays in Xenopus egg extract model to assess the Arpp19-PP2A-B55 interaction and S67/S71 dephosphorylation rate, as well as evaluate the physiologic activity of this inhibitor into promoting mitotic entry.

We first tested the involvement of the KKR residues flanking the DSG motif of Arpp19 in PP2A-B55 binding. These motifs have been proposed to act as PP2A-B55 recognition signals for ENSA and to mediate their interaction. As previously described for ENSA, the mutation of these residues to alanine resulted in a decrease in Arpp19 S71 dephosphorylation rate ^24^. However, in great contrast with this previous study, this decrease was associated with a reinforced, instead of destabilized Arpp19-PP2A-B55 interaction. This data suggests a putative implication of KKR residues in reducing instead of promoting the Arpp19-PP2A-B55 interaction, as previously proposed ^24^. However, our data do not allow us to establish whether these residues modulate Arpp19-PP2A-B55 interaction directly or via increasing phospho-S71 catalysis the two main properties defining Arpp19 inhibition of PP2A-B55.

We next investigated the role of the DSG motif in Arpp19 inhibitory activity. Our data showed that the mutation of the acidic and aromatic residues of this motif into alanine converted Arpp19 into a regular PP2A-B55 substrate. Indeed, we observed an accelerated rate of S71 dephosphorylation thereby conferring to this residue a similar temporal pattern of dephosphorylation than other PP2A-B55 substrates such as Prc1. Due to the proximity of the mutated sites, it is very likely that this acceleration could be at least in part promoted by a modification of the docking of S71 of Arpp19 into the catalytic domain of PP2A-B55. The strict conservation of the DSG motif does not allow the precise prediction of the putative structural role of each of the residues mutated in this study but suggests its important role either for adopting a resting conformation of Arpp19 as an “unfair” substrate or for moving to a dephosphorylation competent conformation. In the latter case, G72 likely allows facile exchange between the two conformations. The large aromatic side-chains of Y68, F69, and Y74 could affect the local conformation prior to or upon binding onto the phosphatase surface. Alternatively, they could also affect the interaction with the phosphatase to favor its binding in the ‘unfair’ conformation. This may involve cross-talks or synergy between these vicinal and hydrophobic aminoacids. The two aspartate residues, D70 and D73, may also regulate the local conformation but an additional role in the observed inhibition could come into play. Indeed, they could also prevent a proper orientation of the phosphate group of S71 by directly interacting with the metallic center of the phosphatase and/or with the two arginines of the PP2A C subunit pointing into the catalytic site (R89 and R214 of human PP2A C subunit) ^28^. Interestingly, this internal competition of D70/D73 with the phosphate of S71 to bind the key catalytic residues of PP2A would be modified in the S71T mutation that would favor a rapid dephosphorylation of this site. Accordingly, upon binding, the whole DSG motif would adopt a constrained conformation, propitious for stable interaction but not for dephosphorylation until slow rearrangement occurs to favorably orient the leaving phosphate group. The distant sites such as the KKR region and the cassette motif could contribute to the optimal “unfair” orientation of the DSG motif upon their docking onto the phosphatase surface. Accordingly, our data support a role of the cassette motif in the modulation of phospho-S71 catalysis without affecting PP2A-B55 binding. Moreover, we proved that a minimal distance is required between this region and the DSG motif. Indeed, when this sequence is shortened, a similar phenotype to those obtained in the DSG mutants was observed. Taking together, we propose a model in which the correct docking of the S71 residue at the PP2A-B55 catalytic site would require the presence of the cassette and the DSG moieties. A folding from the former to the latter would be therefore necessary to acquire the correct spatial constrained phospho-S71 position. This spatial conformation, together with the sequence specificity of the cassette and the DSG motifs, would be essential to confer the slow dephosphorylation of S71 and the “unfair” inhibition of PP2A-B55.

Once the structural properties of Arpp19 were identified, we focused our interest on the regulation of its physiological activity by phosphorylation. Besides Gwl-dependent phospho-S71, phosphorylation of Arpp19 on S113 has also been identified ^17,18^. Notably, it has been shown that this residue is maintained phosphorylated by PKA in prophase I-arrested oocytes and partially drops upon PKA inhibition induced by progesterone ^18^. Moreover, the injection of the S113D phospho-mimetic mutant in these oocytes prevents meiotic resumption by progesterone whereas when a thio-S71-S113 double phosphorylated form is injected, oocytes resume meiosis in the absence of hormone stimulation ^19^. These data suggest a negative modulation of phospho-S113 on S71 phosphorylation, a regulation that could play an essential role in the establishment of the correct temporal pattern of PP2A-B55 inhibition and Cyclin B/Cdk1 activation during oocyte maturation. According to a modulation of S71 phosphorylation by S113, our data demonstrate that phospho-mimetic S113D Arpp19 displays an increased S71 dephosphorylation rate compared to the wildtype form. Interestingly, our NMR analysis point out a chemical shift difference between the wildtype and the S113D mutant for three essential residues of the cassette motif (residues Q109, R110 and K111), suggesting a change into the degree of liberty for those residues. Taking together, this data suggests that S113, when phosphorylated, impacts the capacity of the cassette motif to correctly dock phospho-S71 on the PP2A-B55 catalytic site. As a result, the dephosphorylation of this residue is faster and the rate of the phosphatase inhibition drops.

Considering the important role of the phospho-S113 in regulating Arpp19 phosphorylation at S71 and thus PP2A-B55 inhibition, it was essential to determine the mechanisms controlling the kinetics of this phosphorylation. Although the kinase responsible for S113 phosphorylation has already been identified as PKA, nothing was known about the identity of the counterbalancing phosphatase. Data presented here clearly demonstrate that as for S71, PP2A-B55 itself is the main phosphatase in charge of S113 dephosphorylation. Moreover, we demonstrate that the dephosphorylation of this residue depends on the previous dephosphorylation of S71 and highlights a double feedback loop by which these two residues control the PP2A-B55-mediated dephosphorylation one another. This feedback loop would play an essential role in the maintenance of the correct balance of these two phosphorylations conferring the correct temporal regulation of PP2A-B55 and Cyclin B/Cdk1 activation during oocyte maturation and meiosis exit.

From all these data, it is tempting to hypothesize that during meiosis Arpp19 phosphorylation on S113 by PKA would be required in a first time to prevent premature Arpp19 activation and PP2A-B55 inhibition and to maintain prophase I arrest. Upon progesterone stimulation, PKA is inhibited. This process triggers a drop in S113 phosphorylation, and together with the starter activation of Cyclin B/Cdk1 would increase Gwl-dependent phosphorylation of Arpp19 on S71 in order to inhibit PP2A-B55. However, it is known that phosphorylation of S113 does not fully disappear upon PKA inhibition but remains partially phosphorylated throughout meiosis^18^. Although the exact role of this remaining phospho-S113 is not known, it could contribute in establishing the correct timing of S71 dephosphorylation and as such, in coordinating PP2A-B55 inhibition with Cyclin B/Cdk1 not only at GBVD but also during meiosis I-meiosis II transition. Finally, this phosphorylation could also contribute in the rapid drop of phospho-S71 Arpp19 and PP2A-B55 reactivation at metaphase II exit. This fine-tuned balance would be ensured by a double feedback loop between these two phospho-residues of Arpp19 and its phosphatase PP2A-B55.

In summary, this study established the main structural basis of controlling Arpp19 S71 dephosphorylation and PP2A-B55 inhibitory activity. Additionally, we identified the regulatory mechanism by which S113 modulates this phospho-site. We finally discovered a double feedback loop between these two sites that would coordinate the proper temporal pattern of Arpp19-dependent PP2A-B55 inhibition and Cyclin B/Cdk1 activation during meiotic progression and exit. Further studies will be required to determine whether this tuned regulation of Arpp19 also takes place during mitosis to timely ensure the accurate entry and progression.

## METHODS

### Xenopus egg extracts

Frogs were obtained from « Centre de Ressources Biologiques Xénopes (CRB) of Rennes», France and kept in a *Xenopus* research facility at the CRBM (Facility Centre approved by the French Government. Approval n° B34-172-39). Females were injected with 500 U Chorulon (Human Chorionic Gonadotrophin) and 18 h later laid oocytes were used for experiments. Adult females were exclusively used to obtain eggs. All procedures were approved by the Direction Generale de la Recherche et Innovation, Ministère de L’Enseignement Supérieur de la l’Innovation of France (Approval n° APAFIS#4971-2016041415177715v4).

Kinase-inactivated egg extracts were obtained from laid eggs that were dejellied on 2% cysteine solution pH 7.8, transferred into MMR solution (25 mM NaCl, 0.5 mM KCl, 0.25 MgCl_2_, 0.025 mM NaEGTA, 1.25 mM HEPES-NaOH pH7.7) and washed twice with XB Buffer (50 mM sucrose, 0.1 mM CaCl2, 1 mM MgCl_2_, 100 mM KCl, HEPES pH 7.8). Eggs were subsequently centrifuged twice for 20 minutes at 10 000 g and the cytoplasmic fractions recovered, supplemented with RNAse (10 μg/ml final concentration) and dialyzed versus a solution of 50 mM Tris pH 7.7, 100 mM NaCl overnight to eliminate ATP. Upon dialysis, extracts were ultracentrifuged for 50 minutes at 300 000 g and supernatant recovered for use.

CSF extracts were obtained from dejellied eggs and washed with XB buffer with 5 mM EGTA to prevent activation. Eggs were then centrifuged twice for 20 minutes at 10 000 g and the cytoplasmic fraction recovered for use. mRNAs encoding the constitutive form of rat CamKII comprising amino acids 1–290 were transcribed “*in vitro”* from pCS2-(1-290) rCamK2 plasmid with the SP6 RNA polymerase and translated in reticulocyte lysates. To promote meiotic exit, 2 μl of constitutive CamKII-translated reticulocyte lysates were then supplemented to CSF extracts to promote meiotic exit ^14^.

### Immunoprecipitation/Immunodepletion

Immunoprecipitations/immunodepletions were performed using 10 μl of extracts, 10 μl of protein G-magnetic Dynabeads (Life Technologies), and 2 μg of each antibody. Antibody-linked beads were washed twice with XB buffer, twice with Tris 50 mM, pH 7.5 and incubated for 15 min at room temperature (RT) with 10 μl of *Xenopus* egg extracts. For immunodepletion, the supernatant was recovered and used for subsequent experiments. Four rounds of immunoprecipitation were required to fully deplete endogenous Arpp19 form CSF extracts.

For B55 depletion, 1,5 ml of ATP-devoid extracts were adjusted at a final concentration of 400 mM NaCl and loaded on a 1,5 ml TALON Superflow Metal Affinity Resin column pre-bound with of 2 mg of His-Arpp19 wild type purified protein. The flow through was then collected in different fractions, analyzed by western blot for B55 levels, aliquoted and frozen at −80°c until use.

For immunoblotting, 0.75 μl of egg extract was subjected to SDS-PAGE, transferred to Protran nitrocellulose (Protran, Amersham) or Immobilon membranes (Millipore) and upon blocking using TBST /5% milk or TBST/5% BSA (when phospho-antibodies were used) incubated with the indicated antibodies.

When experiments to assess the capacity of Arpp19 mutants to promote mitotic entry were performed, 0,2 μg of exogenous Arpp19 and 50 ng of K72M Gwl were added in 20 μl of Arpp19-depleted egg extracts and a sample of 2 μl was recovered and used for western blot.

### Plasmids

For (48-120) Arpp19 mutant, a 6His tag was subcloned in the pMal-C2X plasmid in the AscI-FseI cloning site. Arpp19 was subsequently subcloned in this double Maltose Binding Protein (MBP)-6His Tag vector in the BamHI-HindIII site.

Xenopus ENSA was amplified from Xenopus ovary cDNAs and subcloned into the pET15-6His vector in the NdeI-BamHI cloning site.

Human PKA and Prc1 clones were obtained from human ORFeome ^29^ version 8 and subcloned in a pDON gateway vector. cDNAs were subsequently subcloned by Gateway in a pET15b vector.

pCMVsport6–Xenopus B56 gamma was obtained from RZPD Deutsches Ressourcenzentrum für Genomforschung GmbH, amplified by PCR and subcloned at the BamHI-SalI site of pGEX4T2.

### Antibody production

Polyclonal rabbit antibodies against Cter Xenopus Cdk1 peptide NH2-CLDKSSLPANQIR-COOH were used for Cdk1 immunoprecipitation. This peptide was covalently coupled to thyroglobulin using sulfo-MBS for immunization. Polyclonal Abs were then purified using the Cter Cdk1 peptide covalently coupled to BSA-sepharose beads using CNBR-sepharose.

Phospho-S109/S113 Arpp19 antibody was produced using phosphorylated S133 peptide (NH2-CLPQRKP**S(p)**LVASKL-COOH). This peptide was covalently coupled to thyroglobulin protein and injected into one rabbit. Rabbit polyclonal antibodies were then affinity purified against the phosphorylated S113 Arpp19 peptide covalently coupled to BSA-sepharose beads. For anti-Xenopus B56 gamma antibodies, GST-B56 gamma was purified and used to immunize rabbits. Immune serum was first pre-cleared of the anti-GST antibodies in a GST-immobilized column and were subsequently affinity purified on immobilized GST–B56 gamma columns.

### *“In vitro”* phosphorylation

Phosphorylation of Arpp19 on S71 or of ENSA on S67 by Gwl was induced by using GST-K72M hyperactive mutant form of Gwl purified from SF9 cells. For *“in vitro”* phosphorylation reaction, 6His-Arpp19 protein and GST-K72M Gwl kinase were mixed at a final concentration of 0.5 μg/μl and 45 ng/μl respectively in a reaction buffer (1 mM ATP, 2 mM MgCl_2_ and Tris 50 mM).

When thio-phosphorylation was performed, regular ATP was substituted by a final concentration of 1 mM of ATP^γs^. For radio-labeling of 6His-Arpp19 on S71, 6His-Arpp19 and GST-Gwl K72M were mixed at a final concentration of 0.5 μg/μl and 45 ng/μl respectively in a reaction buffer containing 200 μM final ATP concentration and ATPγ^33^P at a specific activity of 5 μCi/ nMol in 2 mM MgCl_2_, 100 mM NaCl and 50 mM Tris pH 7.5.

For phosphorylation of Arpp19 on S113 or ENSA on S109 by PKA, a final concentration of 0.5 μg/μl of Arpp19/ENSA and of 50 ng/μl of 6His-PKA catalytic subunit purified from His-tag column in a reaction buffer containing 1 mM ATP, 2 mM MgCl_2_, 100 mM NaCl and 50 mM Tris pH 7.5 were used.

For phosphorylation of 6His-Prc1 on T481, Cdk1 immunoprecipitate was used. One μg of 6His-human Cyclin A was mixed with 100 μl of CSF extracts during 30 minutes and subsequently supplemented with 50 μl of protein G magnetic Dynabeads pre-linked with 10 μg of Xenopus Cdk1 C-terminus antibodies. After 45 minute-incubation the beads were washed 3 times with 500 mM NaCl, 50 mM Tris pH 7.5, twice with 100 mM NaCl, 50 mM Tris pH 7.5 and finally resuspended with 100 μl of reaction buffer containing 1mM ATP, 2 mM MgCl_2_, 100 mM NaCl and 50 mM Tris pH 7.5. 6His-Prc1 was then added to the beads at a final concentration of 1.5 μg/ul.

All *“in vitro”* phosphorylation reactions were incubated for 1 hour at RT, aliquoted and frozen at −80°C until use.

### Dephosphorylation reactions in ATP-devoid interphase egg extracts

When either S71 or S113 Arpp19, S67 or S109 ENSA or T481 Prc1 dephosphorylation was checked by western blot in ATP-devoid interphase Xenopus egg extracts, 1 μl of the corresponding *“in vitro”* phosphorylation mix was diluted with 9 μl of Tris 50 mM-10 mM EDTA buffer and supplemented with 10 μl of ATP-devoid interphase extracts. Final concentration of the mix was adjusted to 300 mM NaCl with a solution of 5 M NaCl Tris 50 mM pH 7.5 and a sample of 2 μl was recovered at the indicated time-points. From the 2 μl sample, 0,7 μl were used to check Arpp19/Prc1 amounts and the rest to assess phosphorylation of the indicated residues using specific phospho-antibodies. Time-point 0 min was prepared by adding separately 1 μl of extract and of 1 μl of phosphorylated substrate directly in Laemmli buffer. When the impact of Arpp19/ENSA phosphorylation of S71 on S113 Arpp19 was checked in ATP-devoid interphase extracts, 2 μl of *“in vitro”* phosphorylated Arpp19 by PKA sample and 2 μl of the *“in vitro”* phosphorylated Arpp19 by Gwl sample were mixed with 8 μl of Tris 50 mM-10 mM EDTA buffer and 20 μl of ATP-devoid interphase extracts and incubated at room temperature. Four microliters of sample were recovered at the indicated time-points and used for SDS-PAGE and western blot.

For the analysis of the effect of S71 phosphorylation of Arpp19 on T481 Prc1 dephosphorylation, 1 μl of *“in vitro”* phosphorylated Prc1 by Cyclin A/Cdk was mixed with 1 μl of *“in vitro”* phosphorylated Arpp19 by GwlK72M, 8 μl of Tris 50 mM-10 mM EDTA and 10 μl of ATP-devoid interphase extracts and incubated at RT. Two microliters of the mix were recovered at the indicated time-points and used for SDS-PAGE and western blot.

When dephosphorylation of Arpp19 mutants was assessed in ATP-devoid extracts by autoradiography, 1μl of the *“in vitro”* ^33^P labelled GwlK72M phosphorylated Arpp19 mutants were mixed with 10 μl of ATP-devoid extracts an incubated at RT. 2 μl samples were then recovered at the indicated time-points and submitted to western blot and autoradiography. In the case of B55 depleted ATP-devoid interphase extracts, and due to the fact that a residual B55 was occasionally observed in the extracts, 0.5 μg of non-radioactive wild type S71 pre-phosphorylated Arpp19 was supplemented to the reaction volume to ensure a full inactivation of PP2A-B55. Samples were then recovered at the indicated times and submitted to western blot and autoradiography.

Dephosphorylation reactions with purified PP2A-B55 phosphatase were performed as for non-depleted ATP-devoid extracts except that purified preparation of PP2A-B55 was diluted four times with 400 mM Nacl Tris 50 mM pH 7.5.

### Gel filtration

Three millilitres of ATP-devoid interphase extracts were loaded into a Hiload 16/60 Superdex 200 (GE Healthcare, Life Science). Proteins were eluted with a Tris 50 mM pH 7.5, 100 mM NaCl buffer at a constant gradient of 0.9 ml/min into fractions of 1.8 ml. Arpp19 S109/S113 and S67/S71 dephosphorylation activities were determined by mixing 5 μl of each elution fraction and 1 μl of *“in vitro”* phosphorylated Arpp19 sample by either Gwl or PKA. 10- minute incubation for phospho-Arpp19 S113 and 20 minutes for phospho-Arpp19 S71 were used.

### H1 Kinase activity

CSF extract samples (1 μl) were frozen at the indicated times during the experiment. When H1 kinase activity was performed, samples were thawed by the addition of 19 μl of phosphorylation buffer including 2 μCi [γ^33^P] ATP and incubated for 10 minutes at RT. Reactions were stopped by Laemmli sample buffer addition and used for SDS-PAGE and autoradiography.

### Mutagenesis

Deletions and single or double-point mutations of Arpp19 were performed using Pfu ultra II fusion DNA polymerase. Oligonucleotides were purchased from Eurogentec and are detailed in the Supplementary material, Table 1. A pMA-T vector encoding for a DNA in which the DSG domain of Arpp19 sequence was exchanged by the cassette domain (D2-D1 mutant) was synthesized by Geneart (Thermofisher) and sub-cloned into Pet-15 6his vector.

### Protein purification

6His-Xenopus Arpp19 (isoform 1, accession number XM018251008.1) ^14^, 6His-Xenopus ENSA (accession number NM001086605.1), 6His-human Prc1, and 6His-Rat Catalytic Subunit of PKA were produced in *Escherichia coli* and purified using TALON Superflow Metal Affinity Resin. Due to the insolubility of the histidine fusion protein for (48-120) N-terminal deletion mutant of Arpp19, a double tagged MBP-His fusion protein was produced and purified using the histidine tag as reported above.

### PP2A-B55 purification

15 ml of ATP-devoid interphase extracts were supplemented with NaCl to a final concentration of 400 mM NaCl and loaded into a 1 ml TALON Superflow Metal Affinity Resin column pre-bound with 200 μg of His-Arpp19 wild type purified protein. After thoroughly washing with a Tris 50 mM pH 7.5, 400 mM NaCl buffer, elution was performed with 5 ml Imidazole 150 mM in Tris 50 mM pH 7.5, 100 mM NaCl. The eluted fraction was then loaded into a HiLoad 16/60 Superdex 200 column and eluted again with a Tris 50 mM pH 7.5, 100 mM NaCl buffer at a constant gradient of 0.9 ml/min into fractions of 1.8 ml. Fractions from 32 to 42 were pooled, diluted 1/3 in Tris 50 mM pH 7.5 and re-loaded into a MonoQ 5/50GL column GE Healthcare, Life Science) (Supplementary Figure 1A). After washing with a Tris 50 mM pH7.5 buffer, elution was performed with a 0 to 800 mM NaCl linear gradient into fractions of 0.4 ml. After western blot analysis, fractions 15, 16 and 17 were pooled and used for dephosphorylation assays.

### PP2A-B55 binding assays

Twenty microliters of HisPur™NiNTA Magnetic beads (Life Technologies) were washed with XB buffer and supplemented with 1 μg of 6His-wild type or the mutant forms of Arpp19 and incubated at 21°C for 20 min in continuous agitation. Beads were recovered, washed three times with XB buffer and supplemented with 25 μl of CSF extract. Upon 20 min of continuous mixing at 21°C, beads were washed again for three times with RIPA buffer (10 mM NaH2PO4, 100 mM NaCl, 5 mM EDTA, 1% Triton X100, 0.5% deoxycholate, 80 μM β-glycerophosphate, 50 mM NaF, 1 mM DTT), twice with HEPES 50 mM pH 7.4 and supplemented with Laemmli sample buffer and used for western blot with anti-Arpp19 and anti-B55 antibodies.

### NMR analysis

NMR data were recorded using 1 mM uniform [^15^N]-Arpp19 or [^15^N]-S113D-Arpp19 labeled protein in 10 mM Na-Phosphate buffer at pH 6.8, 100 mM NaCl with 10% (v/v) ^2^H2O. Chemical shift assignments were determined from standard three-dimensional NMR experiments (3D [^1^H-^15^N] NOESY-HSQC and 3D [^1^H-^15^N] TOCSY-HSQC using mixing times of 200 and 60 ms, respectively) recorded at 20°C on a 700 MHz Bruker AVANCE spectrometer equipped with a z-gradient ^1^H-^13^C-^15^N triple resonance cryogenic probe. ^1^H chemical shifts were referenced directly, and ^15^N chemical shifts indirectly, to 2,2-dimethyl-2-silapentane-5-sulfonate DSS, methyl proton signal et 0.0 ppm. All NMR spectra were processed and analyzed with Gifa ^30^.

## COMPETING FINANCIAL INTEREST

The authors declare no competing financial interests.

## ACKNOWLEDGMENTS

We are grateful to Marc Plays and Phillipe Richard form animal and antibody production facilities of the CRBM and to Frédéric Lionneton and Sylvie Fromont from Montpellier Genetic Collection facility. We thank Yvan Boublik for help on baculovirus GwlK72M production. We are greatful to Priya Amin for critical reading of this manuscript. This work was supported by the Agence National de la Recherche (KiPARPP, ANR-18-CE13-0013), La Ligue Nationale Contre le Cancer (Equipe Labellisée), the national infrastructure FRISBI (ANR-10-INBS-05) and the LABEX EpiGenMed (ANR-10-LABEX-12-01). P.G.R. has a Labex EpiGenMed and a Fondation de France fellowship. P.R. was granted by “La Ligue National Contre le cancer”.

## AUTHOR CONTRIBUTION

J.C.L., designed and performed biochemical experiments, and analyzed data. S.V. performed biochemical experiments in CSF egg extracts. F.M. helped in the molecular biology experiments. P.R. constructed S113D Arpp19 mutant, purified PKA recombinant protein and performed some Arpp19-rescue experiments. C.G. participated in some Arpp19 rescue experiments. P.G.R. helped on the construction of plasmids and in the production of recombinant proteins. M.G. and P.B. performed NMR experiments and analyzed this part of the result, G.L, using protein modeling, participated in the discussion to provide hypotheses of how the different Arpp19 determinants impact on the oscillation between unfair and fair position of S71P on Arpp19 protein. A.C. designed experiments analyzed data and wrote the paper. T.L., designed experiments, analyzed data, and wrote the paper.

## Notes

### Competing Interest Statement

The authors have declared no competing interest.

## REFERENCES

1. Vigneron, S. et al. Greatwall maintains mitosis through regulation of PP2A. EMBO J 28, 2786–93 (2009).

2. Mochida, S., Ikeo, S., Gannon, J. & Hunt, T. Regulated activity of PP2A-B55 delta is crucial for controlling entry into and exit from mitosis in Xenopus egg extracts. The EMBO journal 28, 2777–85 (2009).

3. Gharbi-Ayachi, A. et al. The substrate of Greatwall kinase, Arpp19, controls mitosis by inhibiting protein phosphatase 2A. Science 330, 1673–7 (2010).

4. Mochida, S., Maslen, S. L., Skehel, M. & Hunt, T. Greatwall phosphorylates an inhibitor of protein phosphatase 2A that is essential for mitosis. Science 330, 1670–3 (2010).

5. Burgess, A. et al. Loss of human Greatwall results in G2 arrest and multiple mitotic defects due to deregulation of the cyclin B-Cdc2/PP2A balance. Proceedings of the National Academy of Sciences of the United States of America 107, 12564–9 (2010).

6. Hached, K. et al. ENSA and ARPP19 differentially control cell cycle progression and development. The Journal of Cell Biology 218, 541–558 (2019).

7. Lindqvist, A., Rodríguez-Bravo, V. & Medema, R. H. The decision to enter mitosis: feedback and redundancy in the mitotic entry network. The Journal of Cell Biology 185, 193–202 (2009).

8. Castilho, P. V., Williams, B. C., Mochida, S., Zhao, Y. & Goldberg, M. L. The M phase kinase Greatwall (Gwl) promotes inactivation of PP2A/B55delta, a phosphatase directed against CDK phosphosites. Molecular biology of the cell 20, 4777–89 (2009).

9. Castro, A. & Lorca, T. Greatwall kinase at a glance. J Cell Sci 131, jcs222364 (2018).

10. Kishimoto, T. Entry into mitosis: a solution to the decades-long enigma of MPF. Chromosoma 124, 417–428 (2015).

11. Crncec, A. & Hochegger, H. Triggering mitosis. FEBS Letters 593, 2868–2888 (2019).

12. Charrasse, S. et al. Ensa controls S-phase length by modulating Treslin levels. Nature communications 8, 206 (2017).

13. Cundell, M. J. et al. The BEG (PP2A-B55/ENSA/Greatwall) pathway ensures cytokinesis follows chromosome separation. Mol Cell 52, 393–405 (2013).

14. Ma, S. et al. Greatwall dephosphorylation and inactivation upon mitotic exit is triggered by PP1. J. Cell. Sci. 129, 1329–1339 (2016).

15. Heim, A., Konietzny, A. & Mayer, T. U. Protein phosphatase 1 is essential for Greatwall inactivation at mitotic exit. EMBO reports 16, 1501–10 (2015).

16. Williams, B. C. et al. Greatwall-phosphorylated Endosulfine is both an inhibitor and a substrate of PP2A-B55 heterotrimers. eLife 3, e01695 (2014).

17. Musante, V. et al. Reciprocal regulation of ARPP-16 by PKA and MAST3 kinases provides a cAMP-regulated switch in protein phosphatase 2A inhibition. Elife 6, (2017).

18. Dupre, A., Daldello, E. M., Nairn, A. C., Jessus, C. & Haccard, O. Phosphorylation of ARPP19 by protein kinase A prevents meiosis resumption in Xenopus oocytes. Nature communications 5, 3318 (2014).

19. Dupré, A.-I., Haccard, O. & Jessus, C. The greatwall kinase is dominant over PKA in controlling the antagonistic function of ARPP19 in *Xenopus* oocytes. Cell Cycle 16, 1440–1452 (2017).

20. Hara, M. et al. Greatwall kinase and cyclin B-Cdk1 are both critical constituents of M-phase-promoting factor. Nat Commun 3, 1059 (2012).

21. Dupre, A. et al. The phosphorylation of ARPP19 by Greatwall renders the auto-amplification of MPF independently of PKA in Xenopus oocytes. Journal of cell science 126, 3916–26 (2013).

22. Chica, N. et al. Nutritional Control of Cell Size by the Greatwall-Endosulfine-PP2A·B55 Pathway. Curr. Biol. 26, 319–330 (2016).

23. Juanes, M. A. et al. Budding yeast greatwall and endosulfines control activity and spatial regulation of PP2A(Cdc55) for timely mitotic progression. PLoS genetics 9, e1003575 (2013).

24. Cundell, M. J. et al. A PP2A-B55 recognition signal controls substrate dephosphorylation kinetics during mitotic exit. The Journal of Cell Biology 214, 539–554 (2016).

25. Hein, J. B., Hertz, E. P. T., Garvanska, D. H., Kruse, T. & Nilsson, J. Distinct kinetics of serine and threonine dephosphorylation are essential for mitosis. Nature Cell Biology 19, 1433–1440 (2017).

26. Dupré, A., Daldello, E. M., Nairn, A. C., Jessus, C. & Haccard, O. Phosphorylation of ARPP19 by protein kinase A prevents meiosis resumption in Xenopus oocytes. Nature Communications 5, (2014).

27. Kumm, E. J. et al. The Cell Cycle Checkpoint System MAST(L)-ENSA/ARPP19-PP2A is Targeted by cAMP/PKA and cGMP/PKG in Anucleate Human Platelets. Cells 9, 472 (2020).

28. Xing, Y. et al. Structure of protein phosphatase 2A core enzyme bound to tumor-inducing toxins. Cell 127, 341–353 (2006).

29. Lamesch, P. et al. hORFeome v3.1: A resource of human open reading frames representing over 10,000 human genes. Genomics 89, 307–315 (2007).

30. Pons, J. L., Malliavin, T. E. & Delsuc, M. A. Gifa V. 4: A complete package for NMR data set processing. J. Biomol. NMR 8, 445–452 (1996).

