## Supplementary Figures and Table for "The study of the determinants controlling Arpp19 phosphatase-inhibitory activity reveals a new Arpp19/PP2A-B55 feedback loop"

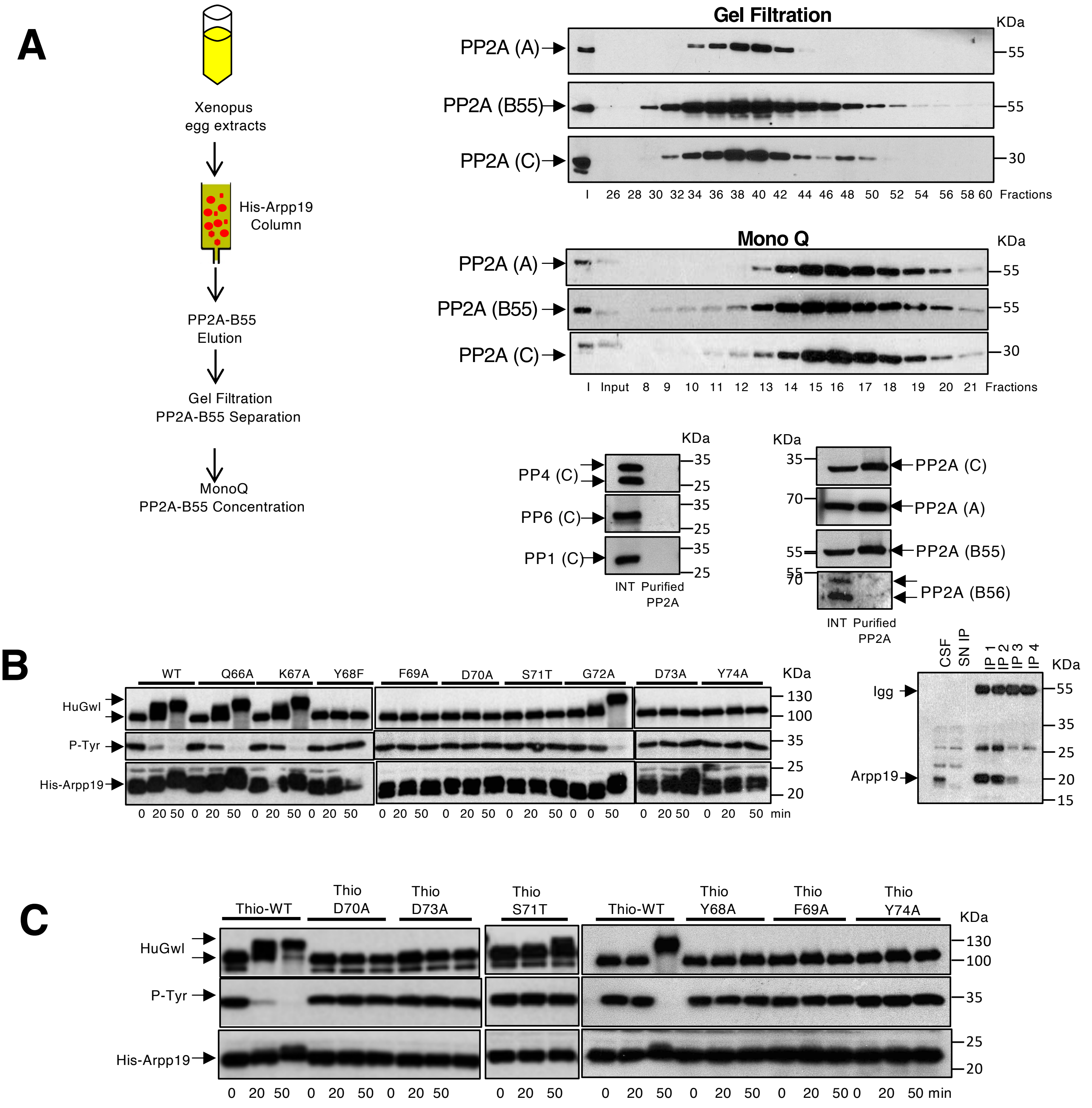

**Supplementary Figure 1.** Aromatic and acidic residues flanking S71 Gwl site of Arpp19 are essential to promote mitotic entry to Arpp19 devoid extracts.

**(A)** Schematic representation of the procedure used to purify PP2A-B55 phosphatase form Xenopus egg extracts. Shown are the amount of PP2A A, B55 and C subunits present in the elution fractions of gel filtration and MonoQ columns. The levels of PP4, PP6, PP1, and A, C, B55 and B56 subunits of PP2A were also examined in the final mix of fractions 15, 16 and 17 representing the pick of purified PP2A-B55 used for the rest of the study.

**(B)** The wildtype or the indicated mutants of Arpp19 were supplemented simultaneously with a trace amount of human GwlK72M and the capacity of this protein to promote mitotic entry in Arpp19-depleted extracts analyzed. The phosphorylation of human ectopic GwlK72M together with the phosphorylation of Tyr 15 of Cdk1 and the levels of ectopic Arpp19 wildtype or mutant proteins were measured by western blot (left panels). In order to fully deplete Arpp19, four rounds of immunoprecipitation were performed in CSF extracts. The amount of Arpp19 remaining upon each immunoprecipitation is shown (right panel). Igg: G Immunoglobulines. SN IP: Supernatant of the fourth immunoprecipitation.

**(C)** As for (B) except that the wildtype or the mutant forms Arpp19 were thio-phosphorylated "*in vitro*" by GwlK72M before being added to the Arpp19-depleted extracts.

Experiments supporting the data of this figure were performed at least three times.

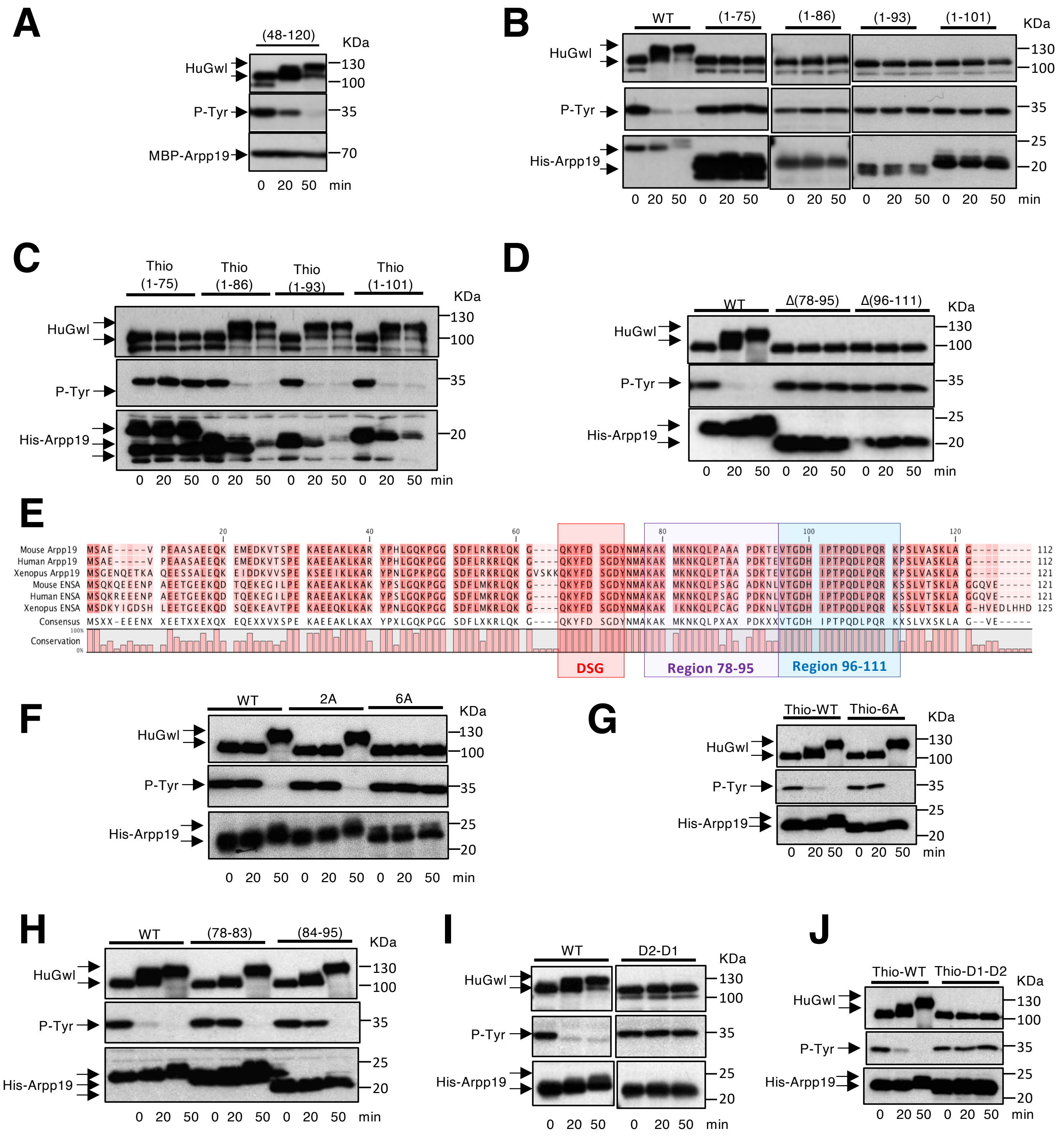

**Supplementary Figure 2.** Arpp19 PP2A-B55 inhibitory activity depends on a specific sequence of the "cassette" motif as well as on a minimal distance between this sequence and the DSG motif.

**(A)** The capacity of the (48-120) Arpp19 mutant to restore mitosis in Arpp19-depleted extracts is determined.

**(B)** and **(D)** as for (A) except that the indicated mutants were tested.

**(E)** Depicted are sequence homology of Arpp19 between different species. Regions 78-95 and 96-11 as well as the DSG motif are indicated with violet, blue and red squares respectively. Note that region 96-111 is highly conserved.

**(F), (H)** and **(I)** as for (A) except that the indicated mutants were analyzed.

**(C), (G)** and **(J)** Rescue assays were performed as for the other mutants except that a thio-S71 phosphorylated form of the indicated mutants was used.

Experiments supporting the data of this figure were performed at least three times.

### KEY RESOURCE TABLE

| REAGENT OR RESOURCE | SOURCE | IDENTIFIER |
|---|---|---|
| **ANTIBODIES** | | |
| Rabbit Polyclonal anti-Human Greatwall | Burgess et al. 2010 | N/A |
| Rabbit Polyclonal anti-Xenopus Cdc27 | Lorca et al. 2010 | N/A |
| Rabbit Polyclonal Phospho-Cdc2 (Tyr-15) | Cell Signaling Technology | Cat#9111 |
| Rabbit Polyclonal anti-Xenopus Cyclin B2 | Abrieu et al. 1997 | N/A |
| Rabbit Monoclonal PhosphoThr320 of PP1 | Abcam | Cat#62334 |
| Mouse Monoclonal Phospho-Erk | Cell Signaling | Cat# 9106S |
| Rabbit Polyclonal anti-Xenopus Arpp19 | Ma et al. 2016 | N/A |
| Rabbit Polyclonal anti-PP2A/B55d | Cell Signaling | Cat#2290 |
| Mouse Monoclonal anti-PP2A (C) subunit, alpha isoform | Merck Millipore | Cat#05-421 |
| Rat Polyclonal PP2A (A) subunit | Cell Signaling | Cat#2260 |
| Rabbit Polyclonal anti-PP2A (B56) gamma subunit | This study | N/A |
| Rabbit Polyclonal anti-PRC1 | Santa Cruz | Cat# 376982 |
| Goat Polyclonal anti-phosphorylated PRC1 (T481) | Santa Cruz | Cat#11768 |
| Mouse Monoclonal anti-PP1 | Kindly gift of Dr M Bollen | N/A |
| Anti-Cter Cdk1 | This study | N/A |
| Rabbit Polyclonal anti-phosphorylated Arpp19 (S67) | Cell Signaling | Cat#11/2017 |
| Rabbit Polyclonal anti-phosphorylated Arpp19 (S109/S113) | This study | N/A |
| Anti-PP4c | Bethyl | Cat#A300-835A |
| Anti-PP6 | Santa Cruz | Cat#393294 |
| Goat anti-rat IgG-HRP | Santa Cruz | Cat#2006 |
| HRP conjugated anti-Rabbit secondary antibodies | Cell Signalling Technology | Cat#7074 |
| Donkey anti-goat IgG-HRP | Santa Cruz | Cat#2020 |
| HRP conjugated anti-Mouse secondary antibodies | BioRad | Cat#1172-1011 |
| **BACTERIAL AND VIRUS STRAINS** | | |
| BL21DE3 Competent E.Coli | New England Biolabs | Cat#C2527H |
| DH5α E. Coli | New England Biolabs | Cat#C2987I |
| **CHEMICALS, PEPTIDES AND RECOMBINANT PROTEINS** | | |
| [gamma-P33] ATP | HARTMANN ANALYTIC | Cat#SRF-301 |
| Pfu ultra II fusion DNA polymerase | Agilent | Cat#600670 |
| $ATP^{\gamma S}$ | Sigma | Cat#A1388 |
| TALON Superflow Metal Affinity Resin | Takara | Cat#635506 |
| PVDF transfer membrane | Millipore | Cat#88518 |
| Protan Nitrocellulose membrane | Amersham | Cat#GE10600016 |
| Dynabeads protein G | Life Technologies | Cat#10004D |
| Histone H1 | Sigma | Cat#14-155 |
| BSA | Sigma | Cat#A7906 |
| His-Pure™NiNTA magnetic beads | Life Technologies | Cat#88832 |
| CNBr-activated sepharose 4B | GE Healthcare | Cat#17-0430-01 |
| Amylose Resin High Flow | Biolabs | Cat#E80225 |
| Sulfo-MBS | Thermo Scientific | Cat#22312 |
| HiLoad™16/600 Superdex | Thermo Scientific | Cat#28-9893-35 |
| MonoQ 5/50 GL | GE-Healthcare | Cat#GE17-5166-01 |
| Recombinant GST-Human Greatwall K72M mutant | Vigneron et al. 2011 | N/A |
| GST-Xenopus B56 gamma | This study | N/A |
| **EXPERIMENTAL MODELS: ORGANISMS/STRAINS** | | |
| Xenopus Laevis | Centre de Ressources Biologiques Xenopes-Rennes | http://www.celphedia.eu/en/centers/crb |
| **OLIGONUCLEOTIDES** | | |
| Forward and reverse primers for Arpp19 K36A/K38A/R40A: | Eurogentec | This study |
| 5’ GAAGTCAGAGGAGATAGCGTTAGCAGCAGCGTATCCTAACCTCGG 3’ | | |
| 5’ CCGAGGTTAGGATACGCTGCTGCTAACGCTATCTCCTCTGACTTC 3 | | |
| Forward and reverse primers for Arpp19 Q66A: | Eurogentec | This study |
| 5’GGCGTAAGTAAAAAGGCAAAATATTTTGACTCTGGG 3’ | | |
| 5’ CCCAGAGTCAAAATATTTTGCCTTTTTACTTACGCC 3’ | | |
| Forward and reverse primers for Arpp19 K67A: | Eurogentec | This study |
| 5’ GGCGTAAGTAAAAAGCAAGCATATTTTGACTCTGGGGAC 3’ | | |
| 5’ GTCCCCAGAGTCAAAATATGCTTGCTTTTTACTTACGCC 3’ | | |
| Forward and reverse primers for Arpp19 Y68A: | Eurogentec | This study |
| 5’ GTAAGTAAAAAGCAAAAAGCTTTTGACTCTGGGGAC 3’ | | |
| 5’ GTCCCCAGAGTCAAAAGCTTTTTGCTTTTTACTTAC 3’ | | |
| Forward and reverse primers for Arpp19 F69A: | Eurogentec | This study |
| 5’ GTAAGTAAAAAGCAAAAATATGCTGACTCTGGGGACTACAT 3’ | | |
| 5’ ATTGTAGTCCCCAGAGTCAGCATATTTTTGCTTTTTACTTC 3’ | | |
| Forward and reverse primers for Arpp19 D70A: | Eurogentec | This study |
| 5’ AAAGCAAAAATATTTTgccTCTGGGGACTACAATATG 3’ | | |
| 5’ CATATTGTAGTCCCCAGAggcAAAATATTTTTGCTTT 3’ | | |
| Forward and reverse primers for Arpp19 S71T: | Eurogentec | This study |
| 5’ GGCCAAAAATATTTTGACACTGGGGACTACAATATGGC 3’ | | |
| 5’ GCCATATTGTAGTCCCCAGTGTCAAAATATTTTTGGCC 3’ | | |
| Forward and reverse primers for Arpp19 G72A: | Eurogentec | This study |
| 5’ CAAAAATATTTTGACTCTGCGGACTACAATATGGCTAAA 3’ | | |
| 5’ TTTAGCCATATTGTAGTCCGCAGAGTCAAAATATTTTTG 3’ | | |
| Forward and reverse primers for Arpp19 D73A: | Eurogentec | This study |
| 5’ TATTTTGACTCTGGGGCCTACAATATGGCTAAA 3’ | | |

| | | |
|---|---|---|
| 5' TTTAGCCATATTGTAGGCCCCAGAGTCAAAATAG 3' | | |
| Forward and reverse primers for Arpp19 Y74A: | Eurogentec | This study |
| 5' TATTTTGACTCTGGGGACGCCAATATGGCTAAAGCA 3' | | |
| 5' TTTAGCCATATTGTAGGCCCCAGAGTCAAAATAG 3' | | |
| Forward and reverse primers for Arpp19 (48-120): | Eurogentec | This study |
| 5' CGCGGATCCAGAAAGCGACTTCAGAAAGGCG 3' | | |
| 5' CCCAAGCTTTCAGCCAGCCAGTTTGCTTGCG 3' | | |
| Forward and reverse primers for Arpp19 (1-75): | Eurogentec | This study |
| 5' GAC-TCT-GGG-GAC-TAC-AAT-TAG-GCT-AAA-GCA-AAG-ATG 3' | | |
| 5' CAT-CTT-TGC-TTT-AGC-CTA-ATT-GTA-GTC-CCC-AGA-GTC 3' | | |
| Forward and reverse primers for Arpp19 (1-86): | Eurogentec | This study |
| 5' GATGAAGAACAAGCAACTGTAAACAGCTGCATCTGATAAA3' | | |
| 5' TTTATCAGATGCAGCTGTTTACAGTTGCTTGTTCTTCATC 3' | | |
| Forward and reverse primers for Arpp19 (1-93): | Eurogentec | This study |
| 5' CAACAGCTGCATCTGATAAATAGGAGGTTACGGGTGATCAT 3' | | |
| 5' ATGATCACCCGTAACCTCCTATTTATCAGATGCAGCTGTTG 3' | | |
| Forward and reverse primers for Arpp19 (1-101): | Eurogentec | This study |
| 5' GTTACGGGTGATCATATTTAGACGCCACAAGACCTCCCT 3' | | |
| 5' GAGGGAGGTCTTGTGGCGTCTAAATATGATCACCCGTAAC 3' | | |
| Forward and reverse primers for Arpp19 D(78-95): | Eurogentec | This study |
| 5' CTCTGGGGACTACAATATGGCTGTTACGGGTGATCATATTCC 3' | | |
| 5' GGAATATGATCACCCGTAACAGCCATATTGTAGTCCCCAGAG 3' | | |
| Forward and reverse primers for Arpp19 D(96-111): | Eurogentec | This study |
| 5' GCTGCATCTGATAAAACGGAGCCGTCTCTCGTTGCAAGCA 3' | | |
| 5' TGCTTGCAACGAGAGACGGCTCCGTTTTATCAGATGCAGC 3' | | |
| Forward and reverse primers for Arpp19 2A: | Eurogentec | This study |
| 5' CGGGTGATCATATTCCTGCGGCACAAGACCTCCCTCAAAG 3' | | |
| 5' CTTTGAGGGAGGTCTTGTGCCGCAGGAATATGATCACCCG 3' | | |
| Forward and reverse primers for Arpp19 6A: | Eurogentec | This study |
| 5' CATATTCCTGCGGCACAAGCCGCCGCTGCAAGGAAACCGTCTCTC 3' | | |
| 5' GAGAGACGGTTTCCTTGCAGCGGCGGCTTGTGCCGCAGGAATATG 3' | | |
| Forward and reverse primers for Arpp19 D(78-83): | Eurogentec | This study |
| 5' TCTGGGGACTACAATATGGCTCAACTGCCAACAGCTGCATCT 3' | | |
| 5' AGATGCAGCTGTTGGCAGTTGAGCCATATTGTAGTCCCCAGA 3' | | |
| Forward and reverse primers for Arpp19 D(84-95): | Eurogentec | This study |
| 5' GGCTAAAGCAAAGATGAAGAACGTTACGGGTGATCATATTCC 3' | | |
| 5' GGAATATGATCACCCGTAACGTTCTTCATCTTTGCTTTAGCC 3' | | |
| Forward and reverse primers for Arpp19 (D2-D1): | Eurogentec | This study |
| 5' GGTCGACAATGTCTGGAGAAAATCAGGAG 3' | | |
| 5' CGCGGATCCTCAGCCTTTAGCCATA 3' | | |
| Forward and reverse primers for Arpp19 S113D: | Eurogentec | This study |
| 5'CTCCCTCAAAGGAAACCGGATCTCGTTGCAAGCAAACTGG3' | | |
| 5'CCAGTTTGCTTGCAACGAGATCCGGTTTCCTTTGAGGGAG 3' | | |
| **RECOMBINANT DNA** | | |
| pCS2-(1-120) rat CamKII | Lorca et al. 1993 | N/A |
| pET7F1-human Cyclin A | Generous gift of G Draetta.; Lorca et al.1992 | N/A |
| pET15-6His-Xenopus Arpp19 | Ma et al .2016 | N/A |
| pFastBac-GST-hGwlK72M | Vigneron et al. 2011 | N/A |
| pET15-6His-Xenopus ENSA | This study | N/A |
| pMalCX2-His-Arpp19 | This study | N/A |
| pET15b-human PKA | This study | N/A |
| pCMVsport6–Xenopus B56 gamma | This study | N/A |
| pET15b-human PRC1 | This study | N/A |
| **SOFTWARE AND ALGORITHMS** | | |
| Adobe Photoshop | Microsoft | Version 12.0x64 |
| Excel | Microsoft | Version 14.7.7 |
| PowerPoint | Microsoft | Version 14.7.7 |

**CONTACT FOR REAGENT AND RESOURCE SHA RING**

**REFERENCES**

mitotic defects due to deregulation of the cyclin B-Cdc2/PP2A balance. *Proceedings of the National Academy of Sciences of the United States of America* **107**, 12564–9 (2010).
op and the Greatwall-PP2A pathway is required for metaphase II arrest and correct entry into the first embryonic cell cycle. *Journal of cell science* **123**, 2281–91 (2010).
does not trigger but delays cyclin degradation in interphase extracts of amphibian eggs. *Journal of cell science* **102 ( Pt 1)**, 55–62 (1992).
ependent protein kinase II mediates inactivation of MPF and CSF upon fertilization of Xenopus eggs. *Nature* **366**, 270–3 (1993).
haracterization of the Mechanisms Controlling Greatwall Activity. *Molecular and Cellular Biology* **31**, 2262–2275 (2011).
all dephosphorylation and inactivation upon mitotic exit is triggered by PP1. *Journal of cell science* **129**, 1329–39 (2016).
e, M. & Picard, A. MAPK inactivation is required for the G2 to M-phase transition of the first mitotic cell cycle. *EMBO J* **16**, 6407–13 (1997).
